# Multiple Dynamical Mechanisms of Phase-2 Early Afterdepolarizations in a Human Ventricular Myocyte Model: *Involvement of spontaneous SR Ca^2+^ release*

**DOI:** 10.1101/613182

**Authors:** Yasutaka Kurata, Kunichika Tsumoto, Kenshi Hayashi, Ichiro Hisatome, Yuhichi Kuda, Mamoru Tanida

## Abstract

Early afterdepolarization (EAD) is known to cause lethal ventricular arrhythmias in long QT syndrome (LQTS). In this study, dynamical mechanisms of EAD formation in human ventricular myocytes (HVMs) were investigated using the mathematical model developed by ten Tusscher & Panfilov (*Am J Physiol Heart Circ Physiol*, 2006). We explored how the rapid (I_Kr_) and slow (I_Ks_) components of delayed-rectifier K^+^ channel currents, L-type Ca^2+^ channel current (I_CaL_), Na^+^/Ca^2+^ exchanger current (I_NCX_), and intracellular Ca^2+^ handling via the sarcoplasmic reticulum (SR) contribute to initiation, termination and modulation of phase-2 EADs during pacing in relation to bifurcation phenomena in non-paced model cells. Dynamical behaviors of the non-paced model cell were determined by calculating stabilities of equilibrium points (EPs) and limit cycles, and bifurcation points. EADs during pacing were reproduced by numerical simulations. Results are summarized as follows: 1) A modified version of the ten Tusscher-Panfilov model with accelerated I_CaL_ inactivation could reproduce bradycardia-related EADs and β-adrenergic stimulation-induced EADs in LQTS. 2) Two types of EADs with different initiation mechanisms, I_CaL_ reactivation–dependent and spontaneous SR Ca^2+^ release–mediated EADs, were detected. 3) Spontaneous SR Ca^2+^ releases occurred at higher Ca^2+^ uptake rates, attributable to the instability of steady-state intracellular Ca^2+^ concentrations. Dynamical mechanisms of EAD formation and termination in the paced model cell are closely related to stability changes (bifurcations) in dynamical behaviors of the non-paced model cell, but they are model-dependent. Nevertheless, the modified ten Tusscher-Panfilov model would be useful for systematically investigating possible dynamical mechanisms of EAD-related arrhythmias in LQTS.

**Key points:**

- We investigated dynamical mechanisms of phase-2 early afterdepolarization (EAD) by bifurcation analyses of the human ventricular myocyte model developed by ten Tusscher and Panfilov.
- A modified version of ten Tusscher-Panfilov model with accelerated inactivation of the L-type Ca^2+^ channel current could reproduce phase-2 EADs in long QT syndrome type 1 and 2 cardiomyocytes.
- Dynamical mechanisms of EAD formation in the paced model cell are closely related to stability and bifurcations of the non-paced model cell.
- EAD mechanisms in the modified ten Tusscher-Panfilov model are different from those in other human ventricular myocyte models in the following respects: 1) EAD formation is partially attributable to spontaneous sarcoplasmic reticulum Ca^2+^ releases; and 2) EAD termination (action potential repolarization) during pacing requires the slowly-activating delayed-rectifier K^+^ channel current.
- The modified ten Tusscher-Panfilov model would be useful for systematically investigating possible dynamical mechanisms of initiation and termination of EAD-related arrhythmias in LQTS.

## Introduction

Early afterdepolarization (EAD) is well known to trigger lethal ventricular arrhythmias, called Torsade de Points (TdP), in patients with long QT syndrome (LQTS) (Weiss *et al*. 2010; Shimizu & Horie, 2011; Shimizu, 2013). For prevention and treatments of ventricular arrhythmias in LQTS patients, therefore, elucidating the mechanisms of initiation and termination of EADs and how to suppress EADs is of crucial importance. There are many experimental studies regarding the mechanisms of EAD formation in cardiomyocytes, suggesting major contribution of reactivation of the L-type Ca^2+^ channel current (I_CaL_) to the initiation of EADs during the action potential (AP) phase 2 (e.g., January *et al*. 1988; January & Riddle, 1989; Guo *et al*. 2007; Weiss *et al*. 2010; Xie *et al*. 2010; Milberg *et al*. 2012; Shimizu, 2013). However, recent experimental studies suggested the major role in EAD formation of the spontaneous Ca^2+^ release from the sarcoplasmic reticulum (SR) (Volders *et al*. 2000; Choi *et al*. 2002; Zhao *et al*. 2012). In our recent theoretical study (Kurata *et al*. 2017) using two human ventricular myocyte (HVM) models developed by Kurata *et al*. (2005) and O’Hara *et al*. (2011), referred to as K05 and O11 models, respectively, we could find EAD formations resulting from the I_CaL_ reactivation, but not the spontaneous SR Ca^2+^ release-mediated EADs. With respect to the termination of EADs (AP repolarization), theoretical studies (Tran *et al*. 2009; Qu *et al*. 2013) using a guinea-pig ventricular myocyte model (Luo & Rudy, 1991) suggested the slowly-activating delayed-rectifier K^+^ channel current (I_Ks_) as a key current to cause termination of EADs. However, our preceding study (Kurata *et al*. 2017) have suggested that the mechanisms of EAD termination are model-dependent, not necessarily requiring I_Ks_. Thus, despite many experimental and theoretical studies, how individual membrane and intracellular components contribute to the initiation, termination and modulations of EADs remains controversial.

The aims of this study were 1) to determine whether the ten Tusscher and Panfilov model (ten Tusscher & Panfilov, 2006; referred to as the TP06 model) for HVMs, which has often been used for simulations of reentrant arrhythmias in the human ventricle (ten Tusscher *et al*. 2004; ten Tusscher & Panfilov, 2006), could reproduce EAD formation in LQTS (validation of the model cell for EAD reproducibility), and 2) to define the contributions of individual sarcolemmal and intracellular components to the initiation, termination, and modulation of phase-2 EADs in the TP06 model in comparison with those in other HVM models (evaluation of model dependence for EAD mechanisms). As in our preceding study (Kurata *et al*. 2017; Tsumoto *et al*. 2017), we examined parameter-dependent changes in stabilities of steady states and AP dynamics in the HVM model from the aspect of bifurcation phenomena, which are parameter-dependent qualitative changes in dynamical behaviors, in nonlinear dynamical systems (Guckenheimer & Holmes, 1983; Parker & Chua, 1989; Kuznetsov, 2003). Conditions and dynamical mechanisms of EAD formation in the paced model cell were determined in relation to bifurcations of the non-paced model cell.

With respect to the dynamical mechanisms of EAD formation, we particularly focused on 1) whether and how contributions of each cellular component to occurrences of EADs and bifurcations in the TP06 model are different from those in the K05 and O11 models; 2) whether spontaneous SR Ca^2+^ release-mediated EAD initiation, which did not occur in the K05 or O11 model, can be reproduced by the TP06 model in connection with a bifurcation (destabilization) of intracellular Ca^2+^ dynamics; and 3) how slow I_Ks_ activation, as well as I_CaL_ inactivation and other slow factors, contributes to EAD termination. This study would further provide a theoretical background for experimental and simulation studies on mechanisms of EAD formation and EAD-triggered reentrant arrhythmias in the LQTS human ventricle, as well as for prevention and treatments of life-threatening arrhythmias, like TdPs, in LQTS.

## Methods

### Mathematical Modeling for HVMs

#### Base mathematical model

In this study, we tested the mid-myocardial (M) cell version of the TP06 model for HVMs (ten Tusscher & Panfilov, 2006), which could reproduce phase-2 EADs during inhibition of I_Ks_ and/or the rapidly-activating delayed rectifier K^+^ channel current (I_Kr_) or enhancement of I_CaL_ when I_CaL_ inactivation was accelerated (Vandersickel *et al*. 2014). The M cell version was chosen because it has smaller I_Kr_ and I_Ks_ and thus more vulnerable to EAD formation than the epicardial or endocardial version, as suggested experimentally as well (Antzelevitch *et al*. 1999). Inconsistent with experimental findings for HVMs (Jost *et al*. 2005; O’Hara & Rudy, 2012), the original version of the TP06 model, which has relatively large I_Ks_ and small I_Kr_, exhibited marked AP duration (APD) prolongation during I_Ks_ inhibition, and failed to reproduce phase-2 EADs during I_Kr_ inhibition. In addition, the Ca^2+^ concentrations in the SR (3–4 mM during 1-Hz pacing) were higher than the experimentally observed values of 1–2 mM for rabbit ventricular myocytes (Shannon *et al*. 2004). Based on these experimental data, therefore, a few modifications were made for the M cell version of the TP06 model. The modified version, referred to as the “mTP06a” model, underwent the following modifications: 1) 60% reduction of the maximum I_Ks_ conductance (g_Ks_) with 50% increment of the maximum I_Kr_ conductance (g_Kr_) to reproduce the I_Kr_/I_Ks_ inhibition experiments, and 40% reduction in the SR Ca^2+^ uptake rate (P_up_) to reduce the Ca^2+^ concentration in the SR during pacing. In a previous theoretical study for EAD formation in the TP06 model (Vandersickel *et al*. 2014), the time constant of I_CaL_ inactivation (τ_fL_) was halved with doubled maximum I_CaL_ conductance (g_CaL_) for facilitating EAD formation. Accordingly, we have further developed another version of the modified TP06 model referred to as the “mTP06b” model with halved τ_fL_ and doubled g_CaL_. **Figure 1** shows simulated behaviors of APs, sarcolemmal ionic currents and intracellular Ca^2+^ concentrations in the original and modified M cell versions of the TP06 model with various g_Ks_ and g_Kr_ values. The modified TP06 models could reproduce the experimentally observed responses of HVMs to reductions of I_Kr_ or I_Ks_, with EADs generated during I_Kr_ reductions only in the mTP06b model. Maximum conductance of the ionic channels, densities of transporters, and SR Ca^2+^ uptake/release rates for the modified versions, as well as for the original version, are given in **Table 1**.

**Table 1:**
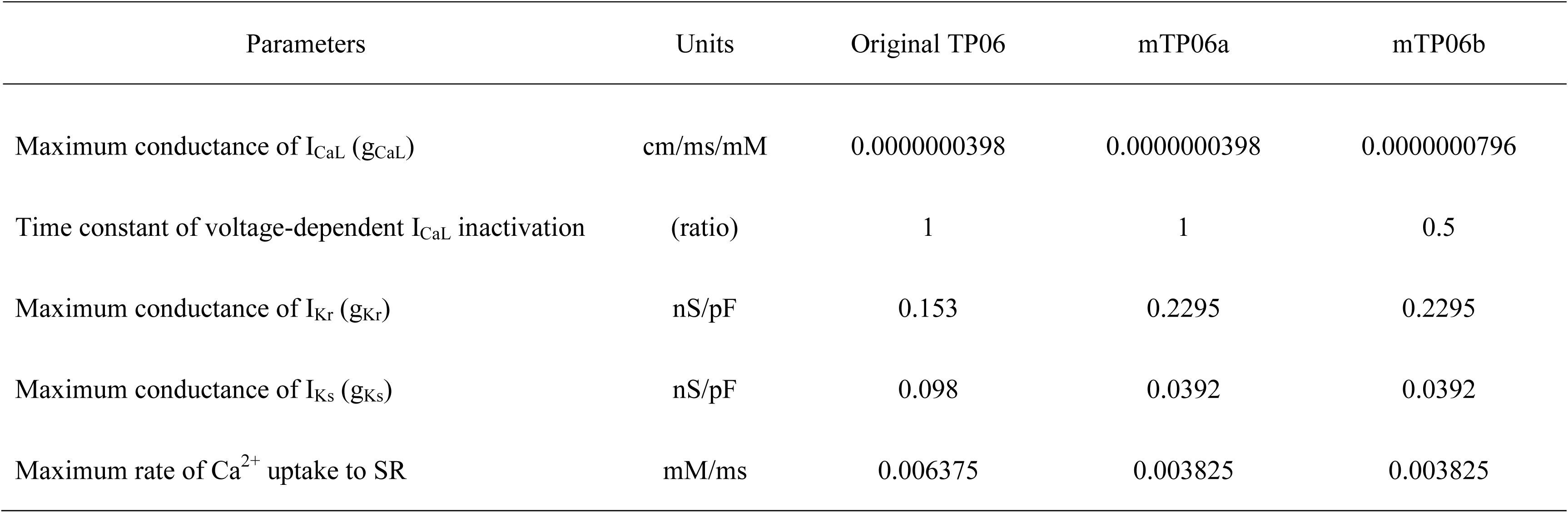
Parameter values for the M cell versions of the mTP06a and mTP06b models relative to those for the original TP06 model.

**Figure 1:**
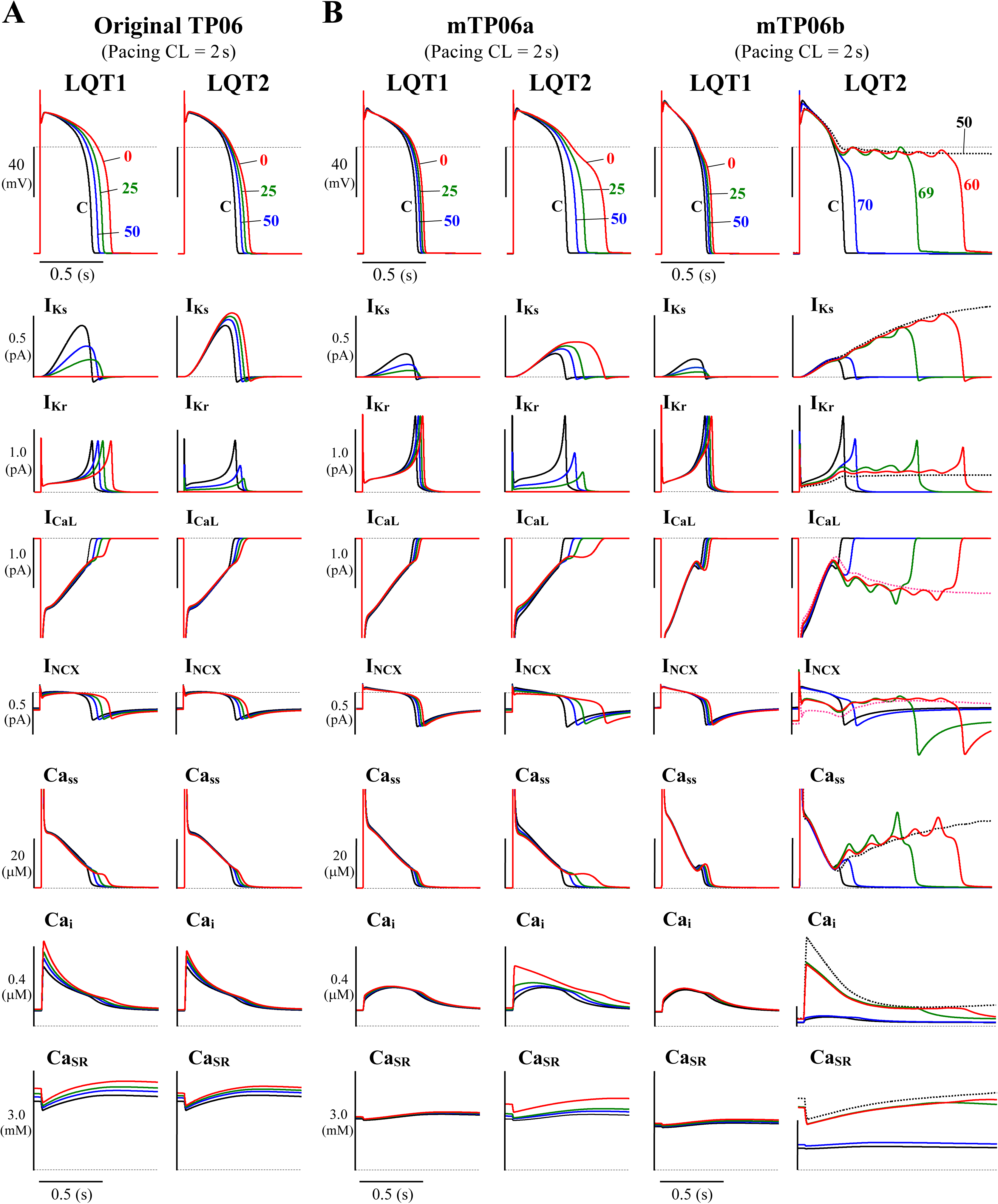
Simulated behaviors of APs (EADs), sarcolemmal ionic currents (I_Ks_, I_Kr_, I_CaL_, I_NCX_) and intracellular Ca^2+^ concentrations (Ca_ss_, Ca_i_, Ca_SR_) in the M cell versions of the original TP06 (A) and mTP06a/b (B) models. To mimic the pathological conditions of LQT1 and LQT2, g_Ks_ and g_Kr_ values, respectively, were decreased by 30–100%; individual APs are labeled by the numbers representing the residual g_Ks_ or g_Kr_ (%Control), with ionic currents and Ca^2+^ concentrations for each g_Ks_ and g_Kr_ value shown by the same colors. The horizontal dashed lines denote the 0 mV, zero current and zero concentration levels. Current amplitudes for the y-axis scale bars are given in pA/pF. The model cells were paced at 0.5 Hz, i.e., with the cycle length (CL) of 2 s for 60 min; AP waveforms, ion currents and Ca^2+^ concentrations after the last stimulus are shown as steady-state behaviors under each condition.

The TP06 model for the normal activity of single HVMs is described as a nonlinear dynamical system of 19 first-order ordinary differential equations. The membrane current system includes the Na^+^ channel current (I_Na_), I_CaL_, I_Kr_, I_Ks_, 4-aminopyridine-sensitive transient outward current (I_to_), inward-rectifier K^+^ channel current (I_K1_), background K^+^ (I_pK_), Na^+^ (I_bNa_) and Ca^2+^ (I_bCa_) currents, Na^+^-K^+^ pump current (I_NaK_), Na^+^/Ca^2+^ exchanger current (I_NCX_), and Ca^2+^ pump current (I_pCa_). Time-dependent changes in the membrane potential (V_m_) are described by the equation,

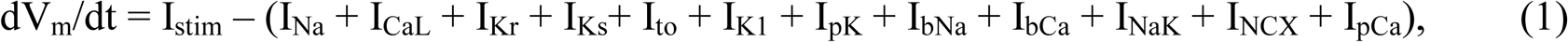

where I_stim_ represents the stimulus current (in pA/pF).

The basic model systems include material balance expressions to define the temporal variations in concentrations of myoplasmic K^+^ (K_i_), Na^+^ (Na_i_) and Ca^2+^ (Ca_i_), and subspace Ca^2+^ (Ca_ss_), while external concentrations of K^+^, Na^+^ and Ca^2+^ were fixed at 5.4, 140 and 2.0 mM, respectively. For bifurcation analyses, K_i_ was fixed at 140 mM for the removal of *degeneracy* (Krogh-Madsen *et al*. 2005; Kurata *et al*. 2008); effects of parameter-dependent changes in K_i_ (∼5 mM) on EAD formation and bifurcation phenomena in the model cell were much smaller than those of the same amount of changes in Na_i_. Na_i_ was unfixed unless otherwise stated, but fixed at 6 mM in some cases (e.g., for the slow-fast decomposition analysis and for voltage-clamped cells, as described later); changes in Na_i_ during AP phase 2 and EAD formation in paced model cells were slow and relatively small.

Details on expressions, standard parameter values, and dynamics of the TP06 model are provided in the original article (ten Tusscher & Panfilov, 2006), and the original TP06 model has been implemented in a cellML-based open resource for public access (http://COR.physiol.ox.ac.uk/). In addition, the original TP06, mTP06a, and mTP06b models have been implemented in PhysioDesigner as XML-based Physiological Hierarchy Markup Language (PHML; http://physiodesigner.org/) models. These models can be referred from PHML database (https://phdb.unit.oist.jp/modeldb/; ID938 to 940), and simulations of their temporal behaviors can be performed using a software, Flint (http://www.physiodesigner.org/simulation/flint/).

#### Modeling LQTS cardiomyocytes with simulated EADs

We developed LQT1- and LQT2-type model cells by simply reducing g_Ks_ and g_Kr_, respectively. As illustrated in **Fig. 1**, the TP06b model, but not the original version (with larger I_Ks_) or the mTP06a model, reproduced phase-2 EADs (and AP repolarization failure) when g_Kr_ became smaller as in LQT2 cardiomyocytes. In contrast, g_Ks_–reduced LQT1 model cells did not exhibit EADs but showed only slight prolongation of APDs under the basal condition, consistent with the recent experimental results from HVMs (Jost *et al*. 2005; O’Hara & Rudy, 2012).

#### Simulating conditions of β-AS

To simulate the condition of β-AS as a major trigger of EADs and TdP in LQT1 patients, we modified the maximum conductance of ion channels and density of transporters based on previous reports (Zeng & Rudy, 1995; Volders *et al*. 2003; Kuzumoto *et al*. 2008), as described in our previous article (Kurata *et al*. 2017). Modifications of parameters for simulating β-AS are listed in **Table 2**.

**Table 2:**
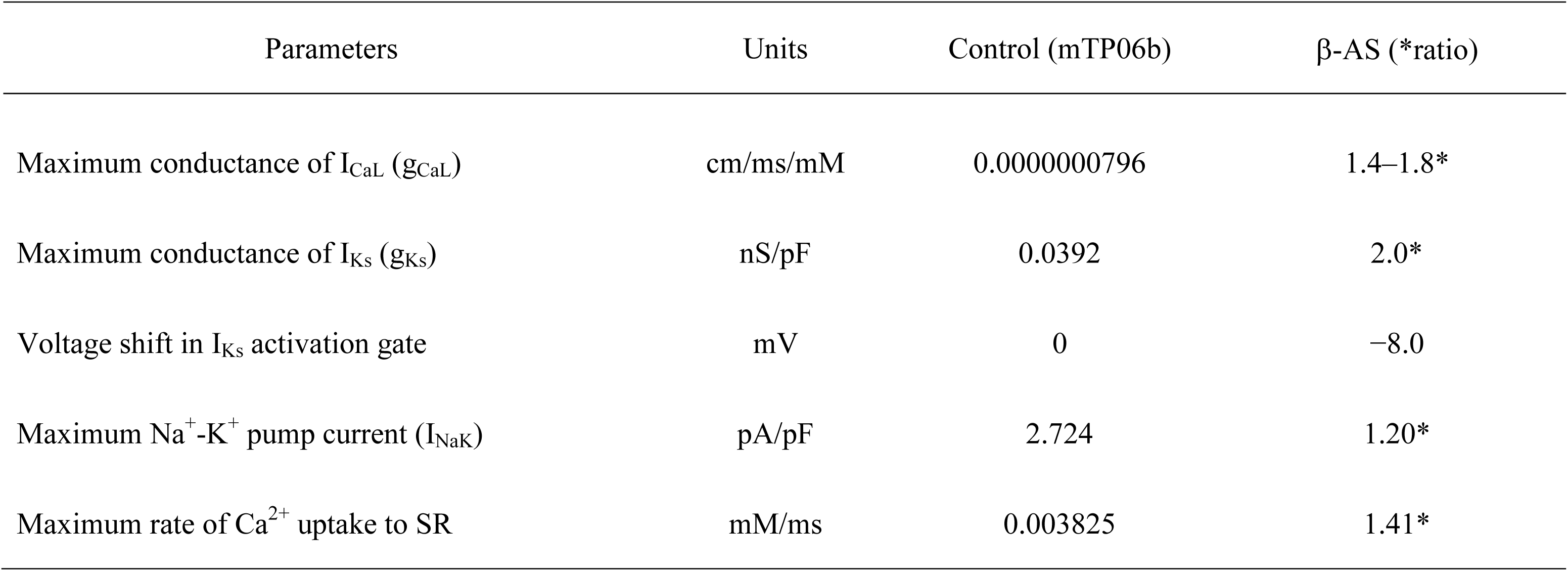
Modifications of parameters for the conditions of β-adrenergic stimulation (β-AS)

### Numerical Methods for Dynamic Simulations

#### Basic methods

Dynamic behaviors of the model cells were determined by numerically solving a set of nonlinear ordinary differential equations including Eq. 1. AP responses were elicited by 1-ms current stimuli of 60 pA/pF at 0.2–2 Hz. Numerical integration was performed by using MATLAB (The MathWorks, Inc., Natick, MA, USA) ODE solvers, *ode 15s* and *ode45*, with the maximum relative error tolerance for the integration methods of 1×10^−8^.

Initial values of the state variables for computation at a parameter set were their steady-state values at a resting V_m_ (see **Table 3** for the control conditions), which were perturbed by the current stimulus; the last values of the state variables in computation were used as initial conditions for the next computation at a new parameter set. The minimum V_m_ during AP phase 4 (V_min_) and the maximum V_m_ during early phase 2 before emergence of an EAD (V_max_), as well as APD at 90% repolarization (APD_90_), were determined for individual APs or AP sets. Steady-state APs for the first parameter set were obtained by numerical integration for 30 min; subsequent numerical integration with each parameter set was continued until the differences in V_min_, V_max_ and APD_90_ between the newly calculated AP and the preceding one became <1×10^−3^ of their preceding values.

**Table 3:**
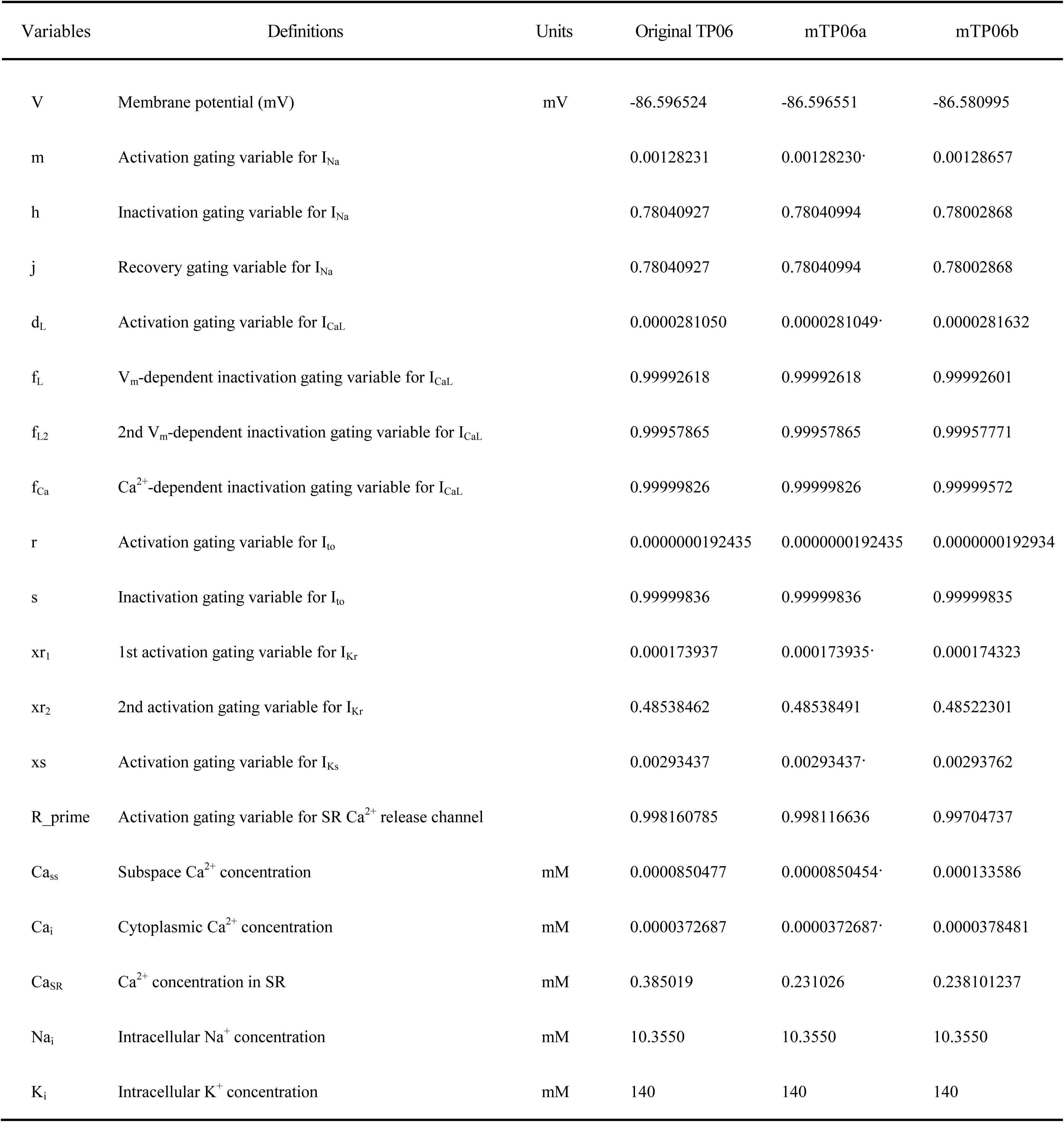
Initial conditions of state variables for computing dynamic behaviors of the M cell versions of the original TP06, mTP06a and mTP06b models as shown in Figure 1.

| Variables | Definitions | Units | Original TP06 | mTP06a | mTP06b |
| --- | --- | --- | --- | --- | --- |
| V | Membrane potential (mV) | mV | -86.596524 | -86.596551 | -86.580995 |
| m | Activation gating variable for $I_{Na}$ | | 0.00128231 | 0.00128230 | 0.00128657 |
| h | Inactivation gating variable for $I_{Na}$ | | 0.78040927 | 0.78040994 | 0.78002868 |
| j | Recovery gating variable for $I_{Na}$ | | 0.78040927 | 0.78040994 | 0.78002868 |
| d<sub>L</sub> | Activation gating variable for $I_{CaL}$ | | 0.0000281050 | 0.0000281049 | 0.0000281632 |
| f<sub>L</sub> | $V_m$-dependent inactivation gating variable for $I_{CaL}$ | | 0.99992618 | 0.99992618 | 0.99992601 |
| f<sub>L2</sub> | 2nd $V_m$-dependent inactivation gating variable for $I_{CaL}$ | | 0.99957865 | 0.99957865 | 0.99957771 |
| f<sub>Ca</sub> | $Ca^{2+}$-dependent inactivation gating variable for $I_{CaL}$ | | 0.99999826 | 0.99999826 | 0.99999572 |
| r | Activation gating variable for $I_{to}$ | | 0.0000000192435 | 0.0000000192435 | 0.0000000192934 |
| s | Inactivation gating variable for $I_{to}$ | | 0.99999836 | 0.99999836 | 0.99999835 |
| xr<sub>1</sub> | 1st activation gating variable for $I_{Kr}$ | | 0.000173937 | 0.000173935 | 0.000174323 |
| xr<sub>2</sub> | 2nd activation gating variable for $I_{Kr}$ | | 0.48538462 | 0.48538491 | 0.48522301 |
| xs | Activation gating variable for $I_{Ks}$ | | 0.00293437 | 0.00293437 | 0.00293762 |
| R_prime | Activation gating variable for SR $Ca^{2+}$ release channel | | 0.998160785 | 0.998116636 | 0.99704737 |
| Ca<sub>ss</sub> | Subspace $Ca^{2+}$ concentration | mM | 0.0000850477 | 0.0000850454 | 0.000133586 |
| Ca<sub>i</sub> | Cytoplasmic $Ca^{2+}$ concentration | mM | 0.0000372687 | 0.0000372687 | 0.0000378481 |
| Ca<sub>SR</sub> | $Ca^{2+}$ concentration in SR | mM | 0.385019 | 0.231026 | 0.238101237 |
| Na<sub>i</sub> | Intracellular $Na^+$ concentration | mM | 10.3550 | 10.3550 | 10.3550 |
| K<sub>i</sub> | Intracellular $K^+$ concentration | mM | 140 | 140 | 140 |

#### Detection of EADs

EADs were detected as transient V_m_ oscillations which emerged during late AP phase 2 (200 ms or later from the AP peak) and eventually led to AP repolarization to a resting V_m_. When phase-2 EADs occurred at higher frequencies of the pacing, a complete AP repolarization was preceded by the next stimulus. Thus, the pacing cycle length (frequency) was usually set to a longer value of 2–5 s (0.2–0.5 Hz), except for analyses of the rate dependence.

All the local minimum (EAD_min_) and maximum (EAD_max_) of V_m_ oscillations during EAD formation, as well as a set of V_min_, V_max_ and APD_90_, were determined for one AP cycle. When APs with EADs were irregular (arrhythmic), all the potential extrema (V_min_, V_max_, EAD_min_, and EAD_max_) and APD_90_ values were sampled for APs evoked by the last 10 stimuli.

### Stability and Bifurcation Analyses for the Non-Paced Model Cell as an Autonomous System

#### Basic methods

We performed bifurcation analysis to explore how dynamical properties of the non-paced autonomous model cell systems alter with changes in parameters. Detailed procedures for bifurcation analyses, i.e., locating equilibrium points (EPs) and limit cycles (LCs), detecting bifurcation points by determination of their stabilities, and constructing one- and two-parameter bifurcation diagrams are provided in our previous articles (Kurata *et al*. 2008, 2012, 2013, 2017; Tsumoto *et al*. 2017). Definitions of the specific terms for bifurcation theory are given in **Table 4**, as well as in textbooks (Guckenheimer & Holmes, 1983; Parker & Chua, 1989; Kuznetsov, 2003). We used 1) a MATLAB nonlinear equation solver (*fsolve*) implementing the Newton-Raphson algorithm to locate EPs and detect bifurcations of EPs; 2) MATLAB ODE solvers (*ode15s* and *ode45*) to calculate stable LCs and detect bifurcations of LCs; and 3) CL_MATCONT, a *continuation toolbox* for MATLAB (Dhooge *et al*. 2006), to locate EPs and LCs as well as to detect bifurcation points. The stability and types of bifurcations of EPs and LCs were determined by calculating eigenvalues of Jacobian matrices for EPs and characteristic multipliers for LCs (Parker & Chua, 1989). As in the K05 and O11 HVM models used for our previous study (Kurata *et al*. 2017), bifurcations to occur in the mTP06 model included 1) Hopf bifurcation (HB) of EPs, 2) saddle-node bifurcation (SNB) of EPs and LCs, 3) period-doubling bifurcation (PDB) of LCs, 4) Neimark-Sacker bifurcation (NSB) of LCs, and 5) *hom*oclinic bifurcation (hom) of LCs (for details, see **Table 4**; also refer to Kuznetsov, 2003; Kurata *et al*. 2017).

**Table 4.** Specific terms for nonlinear dynamics and bifurcation theory.

| Term | Definition |
| --- | --- |
| Autonomous system | An nth-order *autonomous* continuous-time dynamical system is defined by the state equation, $\text{d}\mathbf{x}/\text{d}t = \mathbf{f}(\mathbf{x})$, where the vector field $\mathbf{f}$ does not depend on time but depends only on the state variable $\mathbf{x}$. A dynamical system is *non-autonomous* when the vector field $\mathbf{f}$ explicitly contains time $t$. |
| Equilibrium point (EP) | A time-independent steady-state point at which the vector field vanishes (i.e., $\text{d}\mathbf{x}/\text{d}t = 0$) in the state space of an autonomous system, constructing the steady-state branch in one-parameter bifurcation diagrams. This state point corresponds to the zero-current crossing in the steady-state current-voltage curve. |
| Limit cycle (LC) | An isolated periodic limit set onto which a trajectory is asymptotically attracted in an autonomous system. A stable LC corresponds to an oscillatory state of a cell. |
| Bifurcation | A qualitative change in a solution of ordinary differential equations caused by altering parameters, e.g., a change in the number of EPs or LCs, a change in the stability of an EP or LC, and a transition from a periodic to quiescent state. Bifurcation phenomena that can be observed in cardiac cells include a cessation or generation of pacemaker activity and occurrence of arrhythmic dynamics. |
| Hopf bifurcation | A bifurcation at which the stability of an EP reverses with emergence or disappearance of a LC, occurring when the eigenvalues of a Jacobian matrix for the EP have a single complex conjugate pair and its real part reverses the sign through zero. It is *supercritical* when a stable EP becomes unstable with emergence of a stable LC; it is *subcritical* when an unstable EP becomes stable with emergence of an unstable LC. |
| Saddle-node bifurcation | A bifurcation at which two EPs (steady-state branches) or two LCs (periodic branches) coalesce and disappear (or de novo creation of two EPs or periodic orbits occur). This bifurcation occurs when one of eigenvalues of a Jacobian matrix is zero (EP) or when one of characteristic multipliers becomes $> 1$ (LC). |
| Period-doubling bifurcation | A bifurcation at which a stable period-K solution becomes unstable and spawns a stable period-2K solution, also referred to as a *flip* bifurcation. This occurs when one of characteristic multipliers becomes $< -1$. |
| Neimark-Sacker bifurcation | A bifurcation at which a stable LC becomes unstable with emergence of torus trajectories; it is thus also called a *torus* bifurcation or a Hopf bifurcation on the Poincare section. This bifurcation occurs when a complex conjugate pair of characteristic multipliers comes outside of a unit circle in the complex plane. |
| Homoclinic bifurcation | A bifurcation at which a periodic solution is born from a homoclinic orbit that comes infinitely close to a single equilibrium point at infinity in time. |

#### Construction of one- and two-parameter bifurcation diagrams

In the present study, one- and two-parameter bifurcation diagrams for the non-paced cell model were constructed as functions of bifurcation parameters, including 1) the maximum conductance of ion channel currents (g_Ks_, g_Kr_, and g_CaL_), 2) scaling factor for I_NCX_, 3) P_up_, and 4) pacing cycle length. The maximum conductance of the ionic channel currents and P_up_ are expressed as normalized values, i.e., ratios to the control values. In one-parameter bifurcation diagrams, V_m_ values at EPs (V_E_), potential minimum/maximum of LCs (LC_min_/LC_max_), and bifurcation points were plotted against a given parameter. For construction of two-parameter bifurcation diagrams, the critical values of a first parameter for bifurcations to occur were determined by the one-parameter bifurcation analysis at each second parameter value. The sets of individual bifurcation points of the first parameter were plotted as functions of the second parameter on a two-dimensional parameter plane.

#### Slow-fast decomposition analysis

We further examined the mechanisms of the initiation and termination of EADs by the *slow-fast decomposition analysis*, in which stability and bifurcations of a fast subsystem are determined as functions of a slow variable (Doi *et al*. 2001; Tran *et al*. 2009; Qu *et al*. 2013; Xie *et al*. 2014). This method can define EAD formation as transient trapping of a trajectory of the full system into the quasi-attractor, i.e., quasi-EPs (qEPs) and/or quasi-LCs (qLCs), transiently emerged in the fast subsystem (Qu *et al*. 2013). The gating variable for I_Ks_ activation (***xs***) and Ca^2+^ concentration in the SR (Ca_SR_) were chosen as slow variables; bifurcation diagrams consisting of steady-state branches (for qEPs) and periodic branches (for qLCs) for the fast subsystem were constructed as functions of a slow variable (*xs*^2^ or Ca_SR_), with trajectories of the full system superimposed on the diagrams.

## Results

### Validation and Characterization of the mTP06 models for LQTS HVMs

We first determined whether the mTP06a/b models can mimic the electrophysiological properties of I_Kr_-reduced LQTS type 2 (LQT2) and I_Ks_-reduced LQTS type 1 (LQT1) HVMs, in which EADs occur mainly at lower heart rates (bradycardia), and under β-adrenergic stimulation (β-AS), e.g., during exercise (tachycardia), respectively.

#### Decreases in I_Kr_ and/or I_Ks_ accelerated EAD formation in the mTP06 model

The mTP06b model, but not the original TP06 or mTP06a model, exhibited an AP with EADs when I_Kr_ was inhibited during 0.5-Hz pacing (**Fig. 1**). Similarly, when g_Kr_ was reduced by 40% during 0.2-Hz pacing, we could observe the AP with EADs in the g_Kr_-reduced mTP06b model, as shown in **Fig. 2A-(a)**. This simulated AP with EADs was accompanied by oscillatory reactivation of I_CaL_, and was terminated (i.e., V_m_ went back to the resting V_m_) as I_Ks_ increased (blue arrows for I_Ks_ in Fig. 2A-(a)). Further reducing g_Kr_ by 60% caused another type of EADs with I_Ks_ saturated before applying the second stimulus (**Fig. 2A-(b)**); just before applying this second stimulus, the transient depolarization in plateau phase originating from the spontaneous SR Ca^2+^ release and resulting activation of inward I_NCX_ (red arrows in **Fig. 2A-(b)**). In this case, repolarization did not occur without the next stimulus, i.e., repolarization failure occurred in the non-paced model cell after the cessation of pacing (**Fig. 2A-(c)**).

**Figure 2:**
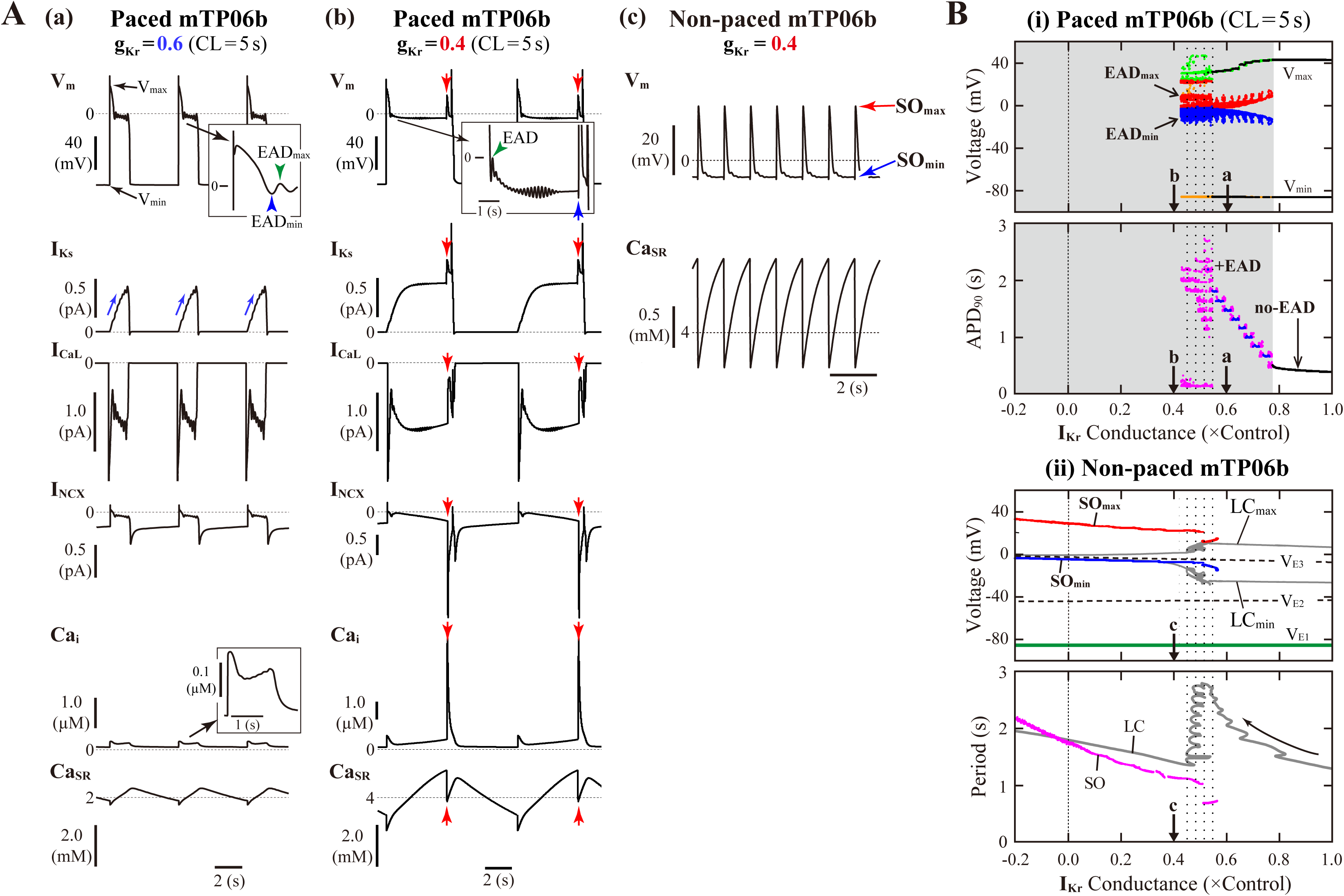
I_Kr_-dependent EAD generation and bifurcations in the mTP06b model. **(A)** Simulated dynamics of V_m_, I_Ks_, I_CaL_, I_NCX_, Ca_i_, and Ca_SR_ (from *top* to *bottom*) in the g_Kr_-reduced model cells paced at 0.2 Hz, illustrating two types of EADs. Temporal behaviors of the variables were computed for 30 min; the behaviors elicited by additional 3–4 stimuli are shown. APs with EADs were terminated (V_m_ repolarized) by gradual increases in I_Ks_ **(a)** or the next stimulus **(b)**. Spontaneous SR Ca^2+^ releases and resulting increases of inward I_NCX_ to provoke EADs occurred at g_Kr_ = 0.4, as indicated by the red arrows (b). With the smaller g_Kr_, sustained V_m_ oscillations driven by spontaneous SR Ca^2+^ releases were observed after cessation of pacing, i.e., in the non-paced model cell **(c)**. **(B)** Potential extrema of APs and EADs, and the AP duration (APD) measured at 90% repolarization (APD_90_) are plotted as functions of normalized g_Kr_ for the paced model cell **(i)**. One-parameter bifurcation diagrams depicting steady-state V_m_ at equilibrium points (EPs), the extrema of limit cycles (LCs) and spontaneous oscillations (SOs), and the periods of LCs and SOs plotted against g_Kr_ are also shown for the non-paced model cell **(ii)**. In the panel **(i)**, AP dynamics during 0.2-Hz pacing were computed for 1 min at each g_Kr_ value, which was reduced from 1.0 to −0.2 at an interval of 0.001. The minimum V_m_ during AP phase 4 (V_min_) and the maximum V_m_ during AP phase 2 before EAD formation (V_max_) are indicated by black dots for rhythmic APs, and by orange (V_min_) and light green (V_max_) dots for arrhythmic APs. When EADs appeared, their local potential minimum (EAD_min_) and maximum (EAD_max_) were plotted by blue and red dots, respectively. In the panel for APD_90_, the black, blue and magenta dots represent APD_90_ values for regular APs without EAD, regular APs with EADs, and arrhythmic APs with EADs, respectively (no-EAD: APs without EAD, +EAD: APs with EADs). The points labeled as “**a**” and “**b**” indicate g_Kr_ values for which AP behaviors are shown in Panel A. In the panel **(ii)**, the steady-state branches as loci of V_m_ at EPs (V_E1-3_), periodic branches as the potential minimum (LC_min_) and maximum (LC_max_) of LCs, and potential extrema of SOs (SO_min_, SO_max_), as well as the periods of LCs and SOs, are plotted for the non-paced model cell. The steady-state branch V_E1_ is stable (green solid lines), while V_E2_ and V_E3_ unstable (black dashed lines). The periodic branches represented by gray solid lines are always unstable. The point labeled as “**c**” indicates the g_Kr_ values for which SOs are shown in Panel A-(c).

**Figure 2B-(i)** shows switching of AP dynamics when g_Kr_ was gradually reduced during 0.2-Hz pacing. EADs emerged at g_Kr_ = 0.772, i.e., with 22.8% block of I_Kr_ (**Fig. 2B-(i)**, *top*); the g_Kr_ reduction led to increases in the number of EADs, resulting in the discrete increase of APD_90_ values (see **Fig. 2B-(i)**, *bottom*). The AP repolarization dynamics in the paced cell model relates to the dynamical behavior of the non-paced cell model because there is no stimulation during the AP repolarization. Therefore, we investigated the dynamical behavior of the non-paced cell model using bifurcation analysis. **Figure 2B-(ii)** shows one-parameter bifurcation diagrams as functions of g_Kr_, constructed for the non-paced mTP06b model (see also **Online Supplementary Fig. S1A** showing those for the mTP06a model for comparison). In the non-paced mTP06b and mTP06a model, there existed three equilibrium potentials (EPs) as the steady states. The EP in the upper steady-state branch (V_E3_) was always unstable at positive g_Kr_ values, while stable at negative g_Kr_ values in the mTP06a model. When g_Kr_ markedly reduced to a large negative value (out of range in **Fig. 2B-(ii)**), the unstable EP (V_E3_) underwent the supercritical HB, which changed it to a stable EP and led to a generation of LC oscillation. The LCs spawned from the HB point were always unstable in the positive g_Kr_ range (see gray lines in **Fig. 2B-(ii)** and **Online Supplementary Fig. S1A-(ii)**). On the one hand, we found small-amplitude spontaneous V_m_ oscillations (SOs) that occurred at depolarized V_m_ (red and blue lines in **Fig. 2B-(ii)**, *top*) in the vicinity of the unstable LCs, as exemplified in **Fig. 2A-(c)**. Just before the disappearance of SOs with increasing g_Kr_, the period of unstable LC markedly prolonged (see the gray zigzag trace in **Fig. 2B-(ii)**, *bottom*). This marked prolongation of LC periods and the emergence of SOs correlated with very long APD (long-lasting EADs) and irregularity of the repolarization time in the paced cell model (compare the dotted ranges in **Figs. 2B-(i)** and **(ii)**).

In contrast, V_E3_ in the I_Ks_-eliminated (g_Ks_ = 0) non-paced model cells was stabilized via an occurrence of the subcritical HB when g_Kr_ was reduced (solid green lines to the left of the label “H” in **Online Supplementary Fig. S1B-(ii)**). Thus, I_Ks_ inhibition caused drastic shift of HB points toward higher g_Kr_ values. Unstable LCs emerged via the subcritical HB, not changing their stability in the g_Kr_ range tested. In the I_Ks_-eliminated paced mTP06a/b models (**Online Supplementary Figs. S1B-(i)**), decreasing g_Kr_ did not yield EADs, but abruptly changed APs without EADs to local responses in the depolarized V_m_ range during pacing, i.e., arrest at stable EPs (V_E3_) without pacing.

To evaluate dependencies of EAD formation on g_Kr_ and g_Ks_, we performed AP simulations using the mTP06b model with various sets of g_Kr_ and g_Ks_. **Figure 3A-(i)** shows a phase diagram of AP behaviors for changes in g_Ks_ and g_Kr_ values with 0.2-Hz pacing. By characterizing AP behaviors observed in the paced mTP06b model, the g_Ks_–g_Kr_ parameter plane was divided into three regions: (1) AP without EAD, (2) AP with EADs (see examples of **Fig. 3B** for the points “**d**” and “**e**” in **Fig. 3A-(i)**), and (3) local response (see an example of **Fig. 3C** for the point “**f**” in **Fig. 3A-(i)**). We further separated the region of the AP with EADs into two regions based on characteristics of the repolarization time in an AP with EADs: During 0.2-Hz pacing, further decreases in g_Kr_ (and/or g_Ks_) in the EAD region altered an AP with shorter APD_90_ of ≤ 5 s that repolarizes before the next stimulus to an AP with longer APD_90_ of > 5 s that is repolarized by the next stimulus, as shown in **Fig. 2A**; then, the APs with EADs were defined as “fast repolarization (fR)” type for the former and “repolarization failure (RF)” type for the latter, which are exemplified in **Fig. 2A-(a)** and **(b)**, respectively. The fR and RF types were distinguished by AP behaviors after an extra stimulus following the last test stimulus to cause AP repolarization, as illustrated in **Fig. 3B**: The fR-type AP repolarized to resting V_m_ within 5 s (**Fig. 3B-(i)**), while the RF-type one did not (**Fig. 3B-(ii)**); in this case, APD_90_ values of the RF-type AP were almost always more than 10 min.

**Figure 3:**
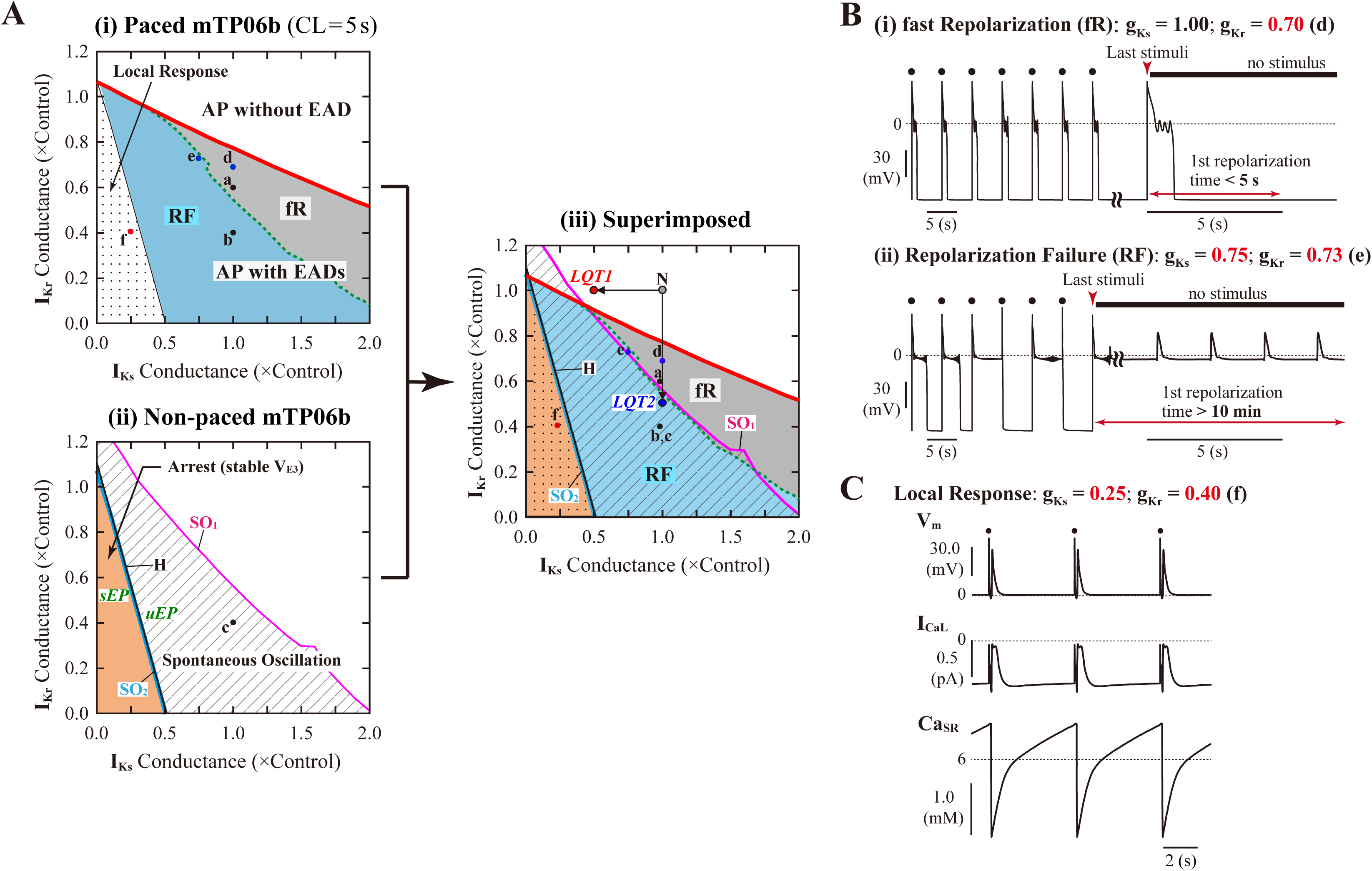
I_Kr_/I_Ks_-dependent EAD formation, dynamics and bifurcations of the mTP06b model. **(A)** A phase diagram indicating the region of EAD formation (and local responses) in the paced model cell **(i)** and two-parameter bifurcation diagrams for the non-paced model cell **(ii)** on the g_Ks_– g_Kr_ parameter plane. In the diagram for the paced model cell (i), the thick red solid, black dashed and thin black solid lines respectively indicate parameter sets of critical points at which short-term EADs (APD_90_ < 5 s, as in Fig. 2A-a and Panel B-(i)), long-term or sustained EADs (APD_90_ > 5 s, as in Fig. 2A-b and Panel B-(ii)), and a local response (Panel C) emerged; parameter regions in which short-term EADs, long-term or sustained EADs, and local responses occur are shown as the light-gray region labeled as “**fR**” (fast repolarization), blue region labeled as “**RF**” (repolarization failure), and dotted region, respectively. In the two-parameter bifurcation diagram for the non-paced model cell (ii), **H**, **SO**_**1**_, and **SO**_**2**_ indicate parameter sets of HB points, critical points at which SOs emerged, and critical points at which SOs switched into quiescence, respectively. The parameter regions in which SOs and convergence to the steady state (V_E3_), i.e., arrest, can be observed are indicated as the shaded and orange regions, respectively. The labels “*sEP*” and “*uEP*” indicate the areas of stable and unstable EPs, respectively, divided by the HB curve. The panel **(iii)** is the diagram for which the phase diagram (i) is superimposed upon the two-parameter bifurcation diagram (ii). The points labeled as “N”, “*LQT1*” and “*LQT2*” denote the normal, LQT1, and LQT2 conditions, respectively. The points “**a**”–“**f**” indicate parameter sets for which AP behaviors are shown in Fig. 2A (a, b, c), and Panels 3B (d, e) and 3C (f). **(B)** Representative behaviors of APs with EADs during 0.2-Hz pacing at the points labeled as “**d**” and “**e**” in Panel A, which are classified into the fast repolarization type (d) and repolarization failure type (e) behaviors, respectively. **(C)** An example of the local response during 0.2-Hz pacing at the point “**f**” in Panel A.

Decreases in g_Kr_ and/or g_Ks_ required for EAD formation were much smaller in the mTP06b model than in the mTP06a model (compare **Fig. 3A-(i)** and **Online Supplementary Fig. S2A-(i)**). The borderline of EAD initiation (the red solid line in **Fig. 3A-(i)**) shifted in a g_Ks_-dependent manner, with the g_Kr_ region of EADs broadening as g_Ks_ increased. Furthermore, two-parameter bifurcation analysis for the non-paced cell model (**Fig. 3A-(ii)**) determined two regions: One is the area of SOs, and the other is the area in which a convergence to depolarized EP (V_E3_), i.e., the arrest at depolarized V_m_ was observed.

To clarify relationships between AP responses observed in the paced cell model and bifurcations occurred in the non-paced cell model, we superimposed the phase diagram on the two-parameter bifurcation diagram (**Fig. 3A-(iii)**). Most of the SO region in which SOs can be observed in the non-paced cell model was included in the RF region, suggesting the relation of spontaneous SR Ca^2+^ release-mediated sustained EADs to SOs (**Fig. 3B-(ii)**). The borderline between local response and AP with EADs corresponded to the HB set in the non-paced cell model, indicating that V_m_ in the paced cell model converges to the stable EP (V_E3_) in the area of local response.

#### Slow pacing facilitated EAD formation in the mTP06b model

To further validate the mTP06b model as a LQT2 model, we next determined whether EAD formation in the g_Kr_-reduced mTP06b model is facilitated at lower pacing rates (in bradycardia). Rate effects on EAD formation are shown in the diagrams depicting the g_Kr_ regions of EADs as functions of the pacing cycle length (**Fig. 4A**). EAD formation in the g_Kr_ reduced system was promoted at lower pacing rates in the Na_i_-variable system, while prevented in the Na_i_-fixed system. As in the K05 and O11 models (Kurata *et al*. 2017), the facilitation of EAD formation at lower pacing rates in the Na_i_-variable mTP06b model was accompanied by the decrease in Na_i_, which resulted in the decrease of outward I_NaK_ leading to delays in AP repolarization and EAD formation (**Fig. 4B-(i)**). In the Na_i_-fixed mTP06b model, the inhibition of EAD formation at lower pacing rates accompanied marked outward shift of I_NCX_ resulting from diminished Ca_i_ transients (**Fig. 4B-(ii)**). In **Online Supplementary Fig. S3**, two-parameter bifurcation diagrams on the g_Ks_–g_Kr_ parameter plane are also shown for the Na_i_-variable and Na_i_-fixed mTP06b model cells paced at 0.2 and 1 Hz. In the Na_i_-variable system (**Online Supplementary Fig. S3A**), slower pacing promoted EAD formation during decreases of g_Kr_ and/or g_Ks_ and broadened the parameter region of EADs; in the Na_i_-fixed system (**Online Supplementary Fig. S3B**), however, the rate-dependent changes in the onset and region of EADs were opposite to those in the Na_i_-variable system (compare the gray and blue areas in each panel of **Online Supplementary Fig. S3**).

**Figure 4:**
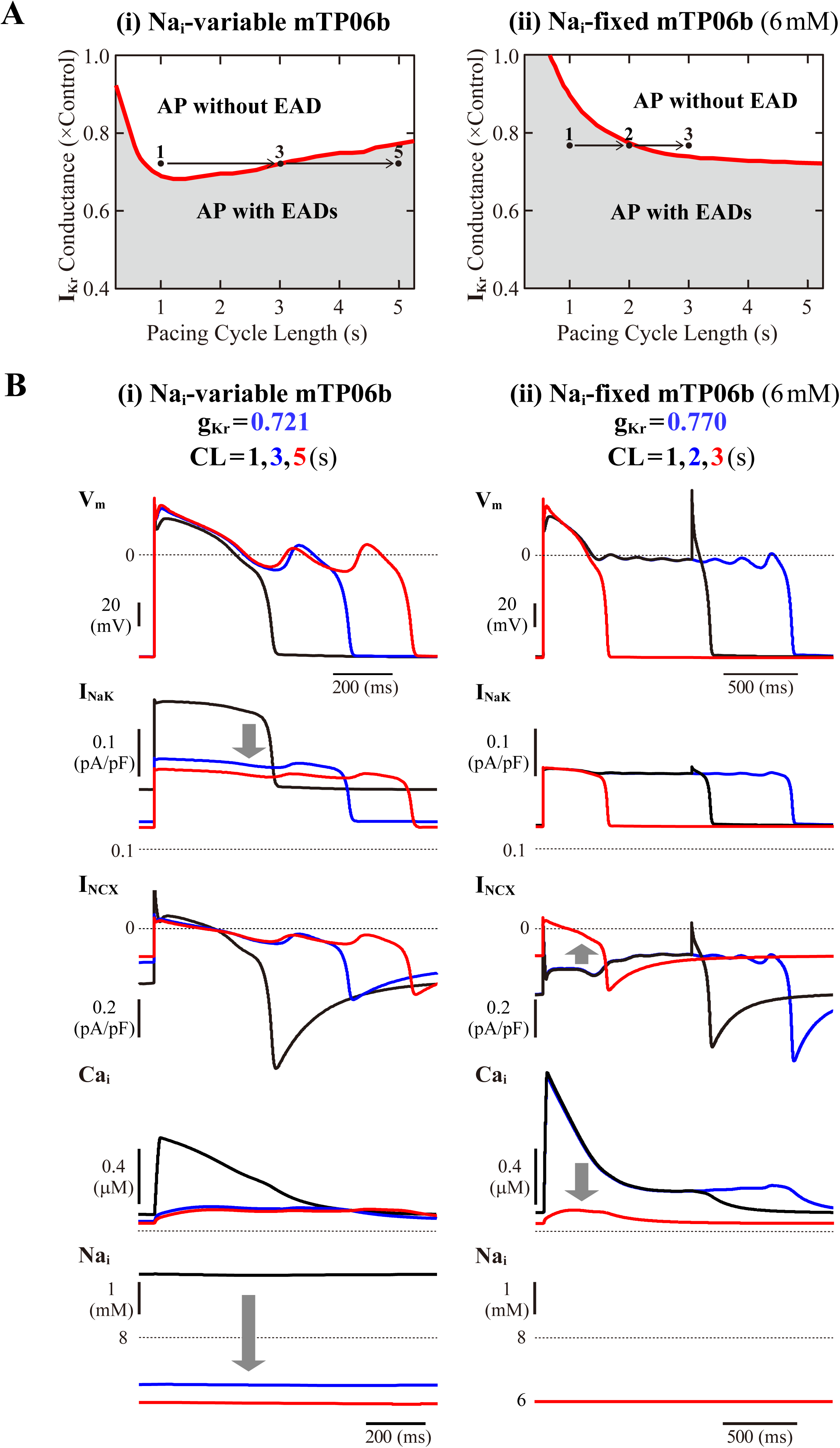
Rate dependence of EAD generation in the mTP06b model. **(A)** Two-parameter phase diagrams for the pacing cycle length (CL) and g_Kr_ depicted for the Na_i_-variable **(i)** and Na_i_-fixed **(ii)** model cells paced with various CLs of 0.75-5.25 s at 0.01-0.2 s intervals. The red solid lines and gray regions represent the parameter sets of critical points for occurrences of EADs and parameter region in which APs with EADs can be ovserved, respectively. **(B)** Simulated dynamics of the Na_i_-variable **(i)** and Na_i_-fixed **(ii)** g_Kr_-reduced model cell paced at various frequencies (with CLs of 1, 3 and 5 s or 1, 2 and 3 s). Temporal behaviors of the model cell were computed for 30 min at each pacing rate; V_m_, I_NaK_, I_NCX_, and Na_i_ for the last 1-2 s are shown as steady-state dynamics.

#### Spontaneous SR Ca^2+^ release-medicated EADs occurred during β-AS

To validate the mTP06b model as a LQT1 model, i.e., to determine whether the I_Ks_-reduced model cell can exhibit EADs under the conditions of β-AS, we examined susceptibilities to EAD generation during β-AS of the normal and LQT1 versions of the mTP06b model. For the LQT1 model cell, g_Ks_ was reduced by 50% and 75%, following the reports for the KCNQ1 mutations M437V and A590W, respectively (see Sogo *et al*. 2016). **Figure 5A** shows simulated APs of the normal and LQT1 versions of the mTP06b model under the basal condition and conditions of β-AS with g_CaL_ increased to 140, 150, 160 and 180% of the control value. The LQT1 model cells exhibited longer APDs under the basal condition (APD_90_ of 334 ms with the normal g_Ks_ vs. 348 ms with 50% g_Ks_ and 356 ms with 25% g_Ks_) and EADs under β-AS with g_CaL_ increased by 60% or more for 50% g_Ks_ and 40% or more for 25% g_Ks_, whereas the normal cell did not exhibit EAD. By constructing a two-parameter bifurcation diagram on the g_Ks_–g_CaL_ plane for the Na_i_-variable model cell paced at 1 Hz (**Fig. 5B**), we could explain their EAD formation under the conditions of β-AS. As in the K05 model (Kurata *et al*. 2017), EAD formation during g_CaL_ increases under β-AS could be inhibited by concomitant g_Ks_ increases more effectively in the normal mTP06b model than in the LQT1 models: The LQT1 model cells entered the area of EAD formation with smaller increases in g_CaL_ (50.8% or more for the 50% g_Ks_ reduction and 31.0% or more for the 75% g_Ks_ reduction), while the normal cell with more than 87.7% increases in g_CaL_. Thus, the mTP06b model could recapitulate EAD formation via enhancement of I_CaL_ during β-AS in the LQT1 cardiomyocyte. Under β-AS with higher g_CaL_ and P_up_, spontaneous SR Ca^2+^ releases as evidenced by abrupt falls in Ca_SR_ without I_CaL_ reactivation often occurred, leading to Ca_i_ elevations (see **Online Supplementary Fig. S4**), increments of inward I_NCX_, and resultant EADs (or spontaneous V_m_ oscillations), as indicated by the dots in **Fig. 5A**.

**Figure 5:**
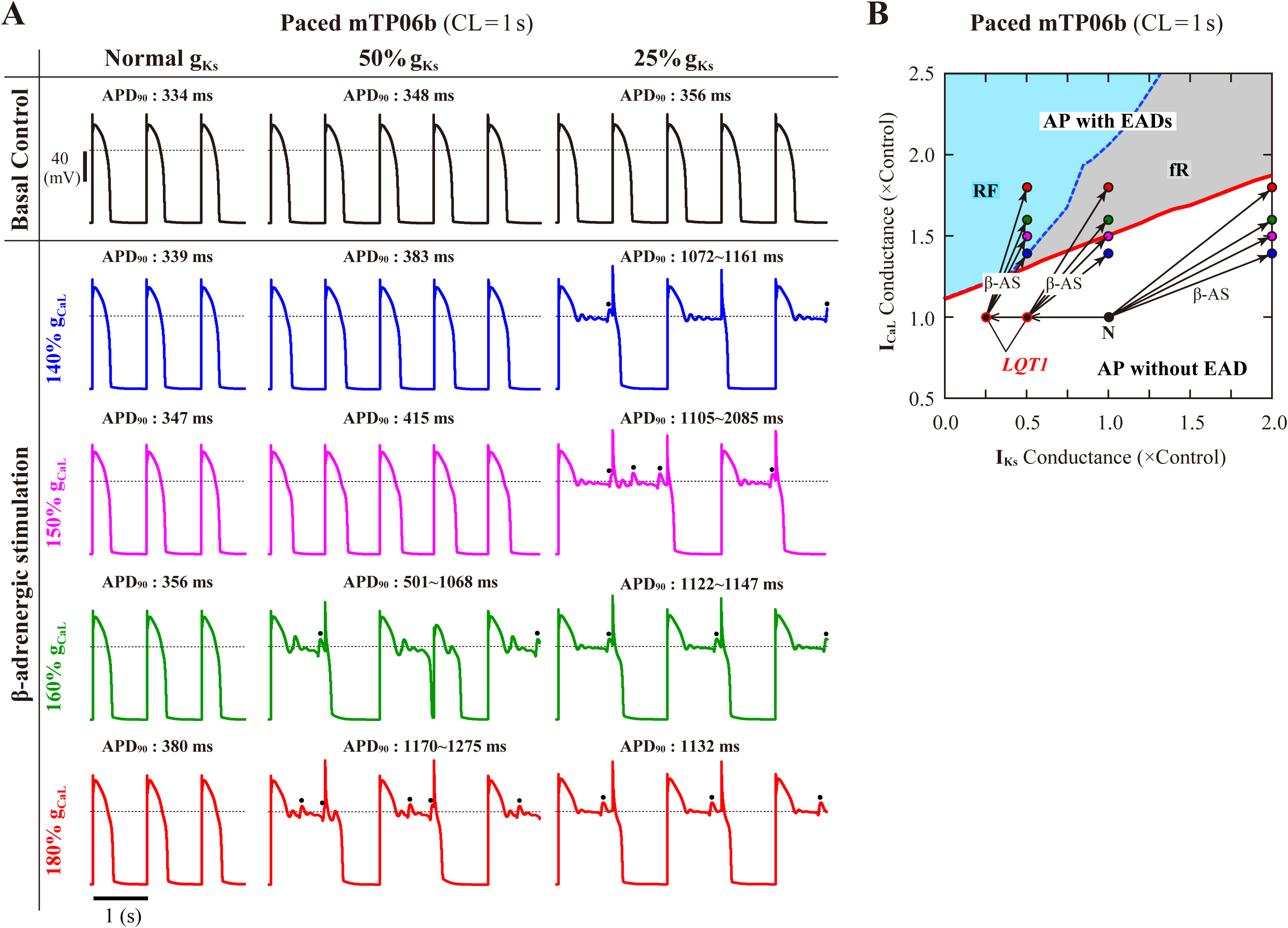
EAD generations during β-AS in the normal and LQT1 versions of the mTP06b model. **(A)**Simulated APs of the model cells under the basal condition (*top*) and conditions of β-AS as indicated by the points and arrows in Panel **B**. Model cells were paced at 1 Hz for 30 min. The dots denote EADs induced by spontaneous SR Ca^2+^ releases. **(B)** A phase diagram on the g_Ks_–g_CaL_ parameter plane depicting displacements of critical points at which EADs emerged during 1-Hz pacing. The critical points were determined during g_CaL_ increases at an interval of 0.002 for individual g_Ks_ values increased at intervals of 0.02–0.1. For simulating the conditions of β-AS, the parameters other than g_CaL_ and g_Ks_ were modified as stated in the Methods section (see Table 2). The *LQT1* model cell was assumed to have reduced g_Ks_ of 50% or 25% of the control value. The points of the control (basal) conditions for cardiomyocytes with the normal and reduced g_Ks_ are labeled as “N” and “LQT1”, respectively. The arrows indicate the parameter shifts from the basal condition to the conditions of β-AS with g_Ks_ doubled and g_CaL_ increased to 140, 150, 160 and 180% of the control value.

#### Influences of SR Ca^2+^ cycling, I_NCX_ and I_CaL_ on EAD formation

Following the finding of spontaneous SR Ca^2+^ releases which occurred especially under β-AS with enhanced I_CaL_ and SR Ca^2+^ uptake, we next examined how SR Ca^2+^ uptake/release (intracellular Ca^2+^ dynamics) and I_NCX_ regulated by the intracellular Ca^2+^, as well as I_CaL_ regulated by the subspace Ca^2+^, affect EAD formation and bifurcations of dynamical behaviors in the mTP06b model by changing P_up_, the scaling factor for I_NCX_, or g_CaL_. **Figure 6A** shows phase diagrams on the P_up_–g_Kr_ and P_up_–g_CaL_ parameter planes. The region of EADs shrank with reducing P_up_ as in the K05 and O11 models (Kurata *et al*. 2017), while broadening at higher P_up_; however, the critical g_Kr_ value (or the critical g_CaL_ value) for the emergence of EADs (red solid lines in **Fig. 6A**) was not decreased (or not increased) but slightly increased (or decreased) as P_up_ reduced. The facilitated EAD formation at smaller P_up_ was associated with increased Ca_i_ and resulting inward shift in I_NCX_ as well as slight increases in I_CaL_ during AP late phase 2 (see **Online Supplementary Fig. S5A**). The two-parameter bifurcation analysis offered further information on how the region of EADs depends on P_up_ and g_Kr_. As shown in **Online Supplementary Fig. S6A**, the critical set of the emergence of EADs and the HB set were mostly parallel to the P_up_ and g_Kr_ axes, respectively, suggesting that alterations in P_up_ contributed not to EAD formation but to rather stability changes of EP (V_E3_) in the non-paced mTP06b model; HB points disappeared with the emergence of spontaneous V_m_ and Ca^2+^ oscillations at higher P_up_, indicating that the SR Ca^2+^ uptake/release machinery destabilizes EPs and thereby induces spontaneous oscillations. When g_Kr_ was markedly reduced in the mTP06b model, spontaneous SR Ca^2+^ releases to cause transient increases in intracellular Ca^2+^ concentrations (Ca_ss_ and Ca_i_) and resulting activation of inward I_NCX_ occasionally occurred with prolonged APD (**Fig. 6B-(a)**). Increasing P_up_ shortened the time to the emergence of the first spontaneous Ca^2+^ release and raised the incidence and frequency of spontaneous Ca^2+^ oscillations to yield EADs or V_m_ oscillations (**Fig. 6B-(b)**).

**Figure 6:**
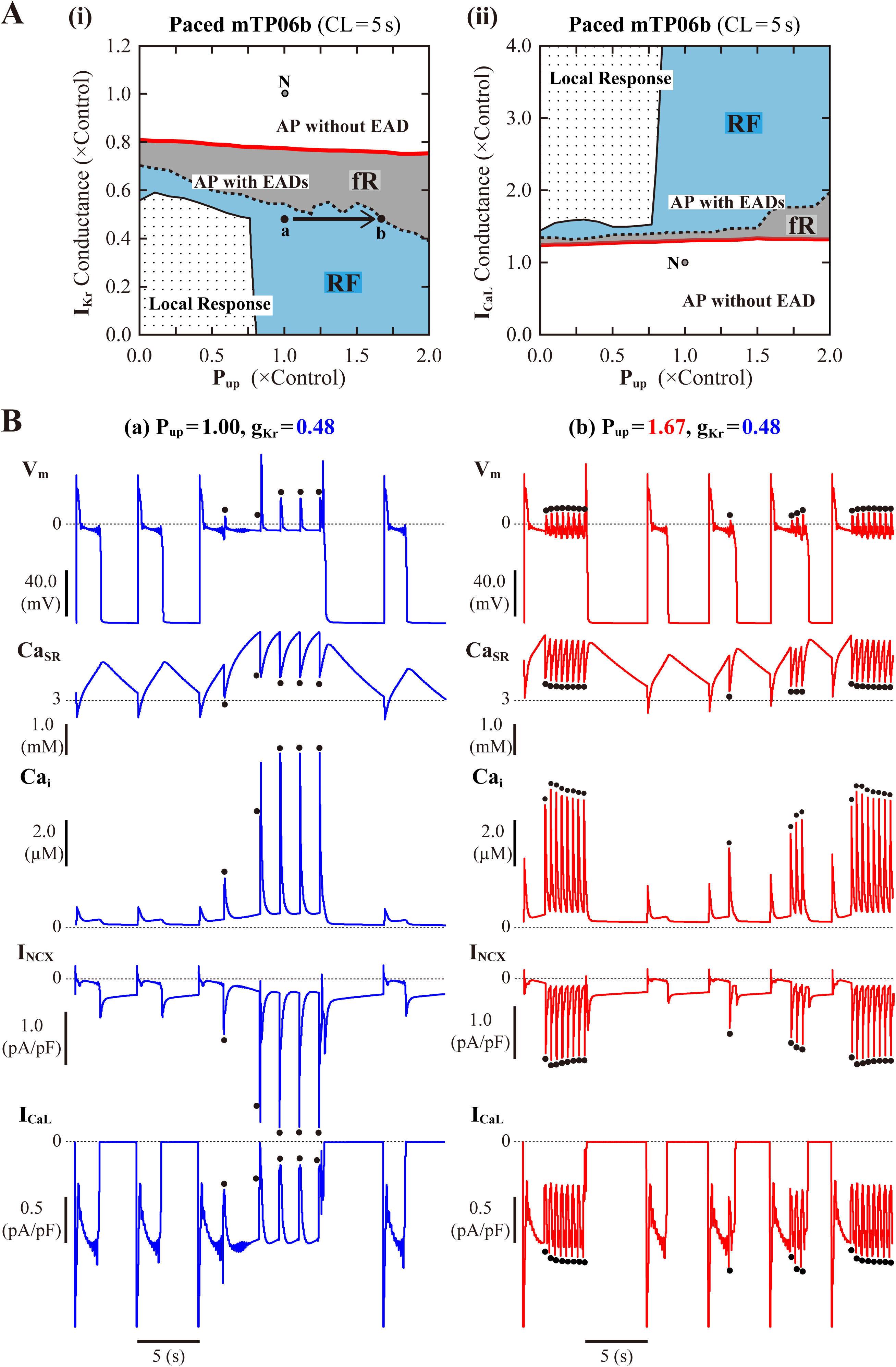
Influences of SR Ca^2+^ handling on EAD generation in the mTP06b model. **(A)** Phase diagrams on the P_up_–g_Kr_ **(i)** and P_up_–g_CaL_ **(ii)** parameter planes, depicting displacements of critical points for the occurrence of short-term EADs (red solid lines) and long-term or sustained EADs (dashed lines) as well as the emergence of local responses (black solid lines). By the parameter sets of these critical points, the parameter planes are divided into the areas of APs with short-term EADs (fR), APs with long-term or sustained EADs (RF) and local response, as described for Fig. 3A. The points “**a**” and “**b**” in the panel (i) denote the parameter sets for simulations of the model cell dynamics shown in Panel B-(a) and (b), respectively. **(B)** Simulated dynamics of the model cells with the normal (1.00) or increased (1.67) P_up_ and the reduced g_Kr_ (0.48). Temporal behaviors of the model cells were computed for 30 min with pacing at 0.2 Hz; V_m_, Ca_SR_, Ca_i_, I_NCX_ and I_CaL_ for additional 30 s are shown as steady-state dynamics. The dots indicate spontaneous SR Ca^2+^ releases as evidenced by abrupt falls of Ca_SR_ and resulting increases in Ca_i_ and inward I_NCX_.

Contributions of I_NCX_ to bifurcations and EAD formation were also explored in relation to those of intracellular Ca^2+^ dynamics, SR Ca^2+^ cycling, and I_CaL_. On the I_NCX_–g_Kr_ and I_NCX_–g_CaL_ parameter planes (**Online Supplementary Fig. S7**), enhancement of I_NCX_ yielded the upward shift in the critical g_Kr_ and downward shift in the critical g_CaL_ for EAD formation (see red curves in **Online Supplementary Fig. S7**), consistent with our previous report for the K05 and O11 models (Kurata *et al*. 2017). Unlike their HVM models, however, the TP06b model did not exhibit a significant shift in the critical g_Kr_ or g_CaL_ for EAD formation when I_NCX_ was reduced; only small inhibition of I_NCX_ was effective in shifting the critical points toward the prevention of EADs. Whether EADs emerge or not depended mainly on the amplitude of inward I_NCX_ and I_CaL_ during the AP late phase 2: As exemplified in **Online Supplementary Fig. S5B**, disappearance of EADs with lower I_NCX_ density was accompanied by a decrease of inward I_NCX_ and a slight reduction of I_CaL_ with increased inactivation during the preconditioning phase just before initiation of the first EAD (see the ellipses and inset in **Online Supplementary Fig. S5B**).

We finally examined effects of I_CaL_ on EAD formation in the paced mTP06b model and bifurcations of dynamical behaviors in the non-paced mTP06b model by changing g_CaL_. g_CaL_-dependent changes in AP dynamics observed in the paced model cell when I_Ks_ was normal (g_Ks_ = 1) and one-parameter bifurcation diagrams as functions of g_CaL_ for the non-paced cell model are shown in **Figs. 7A-(i)** and **7A-(ii)**, respectively. The one-parameter bifurcation diagrams for g_CaL_ (**Fig. 7A-(ii)**) suggest the scenario of EAD formation during enhancement of I_CaL_, which is different from those in the K05 and O11 models (Kurata *et al*. 2017): Increments of g_CaL_ yielded unstable EPs via a SNB of EPs and unstable LCs via a SNB of LCs. With normal I_Ks_ (g_Ks_ = 1), an enhanced g_CaL_ of 1.298-fold the control value was high enough for EAD formation in the mTP06b model (**Fig. 7A-(i)**), whereas unrealistically large increases in g_CaL_ (to 4.248-fold the control value) were required in the mTP06a model (**Online Supplementary Fig. S8A**). **Figure 7B** shows a phase diagram of AP behaviors in the paced model cell (**Fig. 7B-(i)**) and a two-parameter bifurcation diagram for the non-paced model cell (**Fig. 7B-(ii)**), as well as the merged diagram (**Fig. 7B-(iii)**), on the g_CaL_–g_Ks_ parameter planes. Decreasing g_Ks_ shifted the critical g_CaL_ value for EAD generation toward lower values and enlarged the g_CaL_ region of EADs (RF) in the mTP06b model. Larger g_CaL_ (going into the RF region in **Fig. 7B-(i)** and **(iii)**) led to the AP behavior classified into the RF type with small-amplitude spontaneous V_m_ oscillations around unstable LCs. EPs (V_E3_) in the mTP06 models were unstable independent of g_CaL_ unless g_Ks_ was extremely low or high; no HB occurred for moderate variation of g_Ks_ value and consequently stability changes of the EP did not occur (see also **Fig. 7A-(ii)**). In the I_Ks_-eliminated mTP06a/b models (g_Ks_ = 0), an EP (V_E3_) was stabilized via supercritical HBs at relatively small g_CaL_; stable LCs emerging from the HBs were immediately destabilized via a PDB or NSB (**Online Supplementary Fig. S8B**). EAD did not occur at g_Ks_ = 0; larger I_CaL_ caused repolarization failure, in this case, local response.

**Figure 7:**
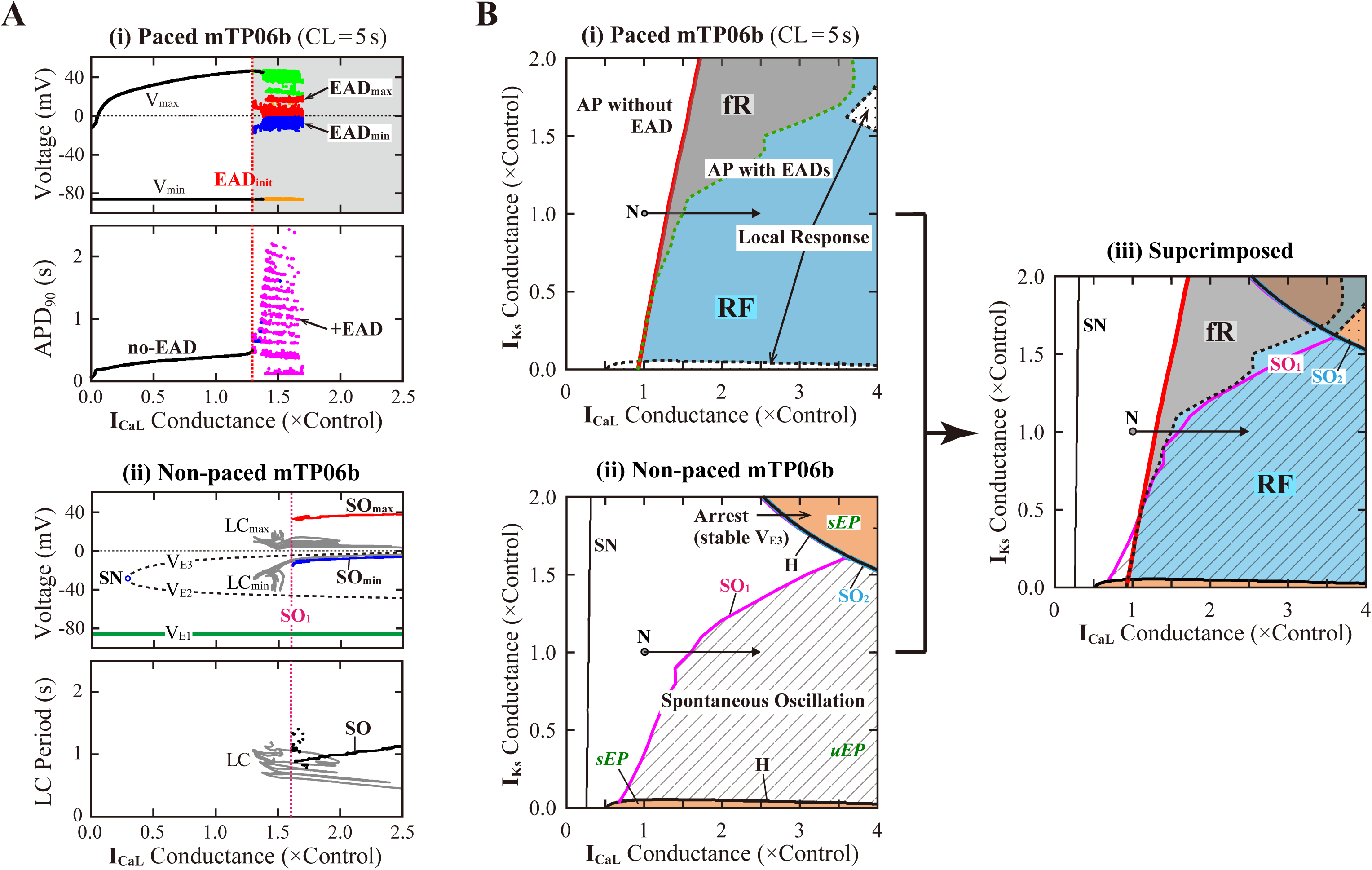
I_CaL_–dependent EAD generation and bifurcations in the mTP06b models. **(A)** Potential extrema of simulated AP and EAD behaviors and APD_90_ for the paced model cell **(i)**, as well as one-parameter bifurcation diagrams with steady-state V_m_ (V_E1–3_), potential extrema of LCs and SOs, and the periods of LCs and SOs for the non-paced model cell **(ii)**, plotted as functions of normalized g_CaL_. Representations and symbols are the same as in **Fig. 2B**. **(B)** A phase diagram **(i)** for the paced model cell, a two-parameter bifurcation diagram **(ii)** for the non-paced model cell, and the merged diagram for which the phase diagram is superimposed upon the two-parameter bifurcation diagram on the g_CaL_–g_Ks_ parameter plane, depicting displacements of critical points at which EADs and local responses emerged, HB points (H), SNB points where unstable V_E2_ and V_E3_ coalesced and vanished (SN), critical points at which SOs emerged (SO_1_) or switched into quiescence (SO_2_), and critical points at which repolarization failure occurred. In the panel **(i)**, the g_CaL_–g_Ks_ parameter plane is divided into the areas of APs without EAD, APs with EADs repolarization of which occurred within 5 s after the last stimulus (fR), and repolarization failure (RF), as in Fig. 3A-(i). The points labeled as “N” denote the normal condition. In the panel **(ii)**, SOs appeared in the shaded region surrounded by SO_1_, SO_2_ and H curves.

### Dynamical Mechanisms for Initiation and Termination of EADs in the mTP06 Model

#### I_Ks_ activation-dependent bifurcations of the fast subsystem associated with EAD formation

To clarify the dynamical mechanisms of EAD formation in the I_Kr_-reduced LQT2-type mTP06b model and why EADs emerge at larger g_Kr_ (or smaller g_CaL_) in the mTP06b model than in the mTP06a model (**Figs. 2** and **7** vs. **Online Supplementary Figs. S1** and **S2**), we further performed the *slow-fast decomposition analysis* (Doi *et al*. 2001; Xie *et al*. 2014). The I_Ks_ activation gating variable ***xs*** or I_Ks_ channel open probability (***xs***^2^) appears to be a slow variable yielding the termination of EADs (**Fig. 2A-(a)**, *the second from top*). Thus, bifurcation diagrams for the fast subsystem composed of the state variables other than the slow variables ***xs***, Na_i_ and Ca_SR_ were first constructed as functions of ***xs***^**2**^, with Na_i_ and Ca_SR_ fixed at constant values (**Fig. 8**, *left*); then, trajectories of the full system (with fixed Na_i_ and Ca_SR_) were superimposed on the diagrams (**Fig. 8**, *right*). The qEP, defined as a steady state of the fast subsystem, at depolarized quasi-V_m_ (qV_E3_) has possessed a property of spiral sink in the mTP06b model (**Fig. 8A**) but spiral source in the mTP06a model (**Fig. 8B**) at ***xs***^2^ = 0. Stable qEP in the former was destabilized via an HB as ***xs***^2^ increased. The g_Kr_ reduction led to broadening of the ***xs***^**2**^ region of stable qEPs (compare green traces of qV_E3_ in **Fig. 8A-(i)** and **(ii)**, *left*). The g_Kr_ reduction-induced broadening of the ***xs***^2^ range of stable qEPs yielded a transient trapping of the full system trajectory in the attractor basin of the stable qEP (**Fig. 8-(ii)**, *right*). This trapping of the full system trajectory around the stable qEP as spiral sink sustained until the trajectory came across the steady-state ***xs***^2^ curve. This trapping phenomenon was not observed in the mTP06a model (**Fig. 8B**, *right*) or the I_Kr_-normal mTP06b model (**Fig. 8A-(i)**, *right*), because the full system trajectories did not intersect with the stable steady-state branch (qV_E3_) before intersecting the steady-state ***xs***^2^ curve. These results indicate that an acceleration of the voltage-dependent I_CaL_ inactivation to form the mTP06b model from the mTP06a model plays a critical role in the stabilization of qV_E3_, consequently leading to the trapping of the full system trajectory.

**Figure 8:**
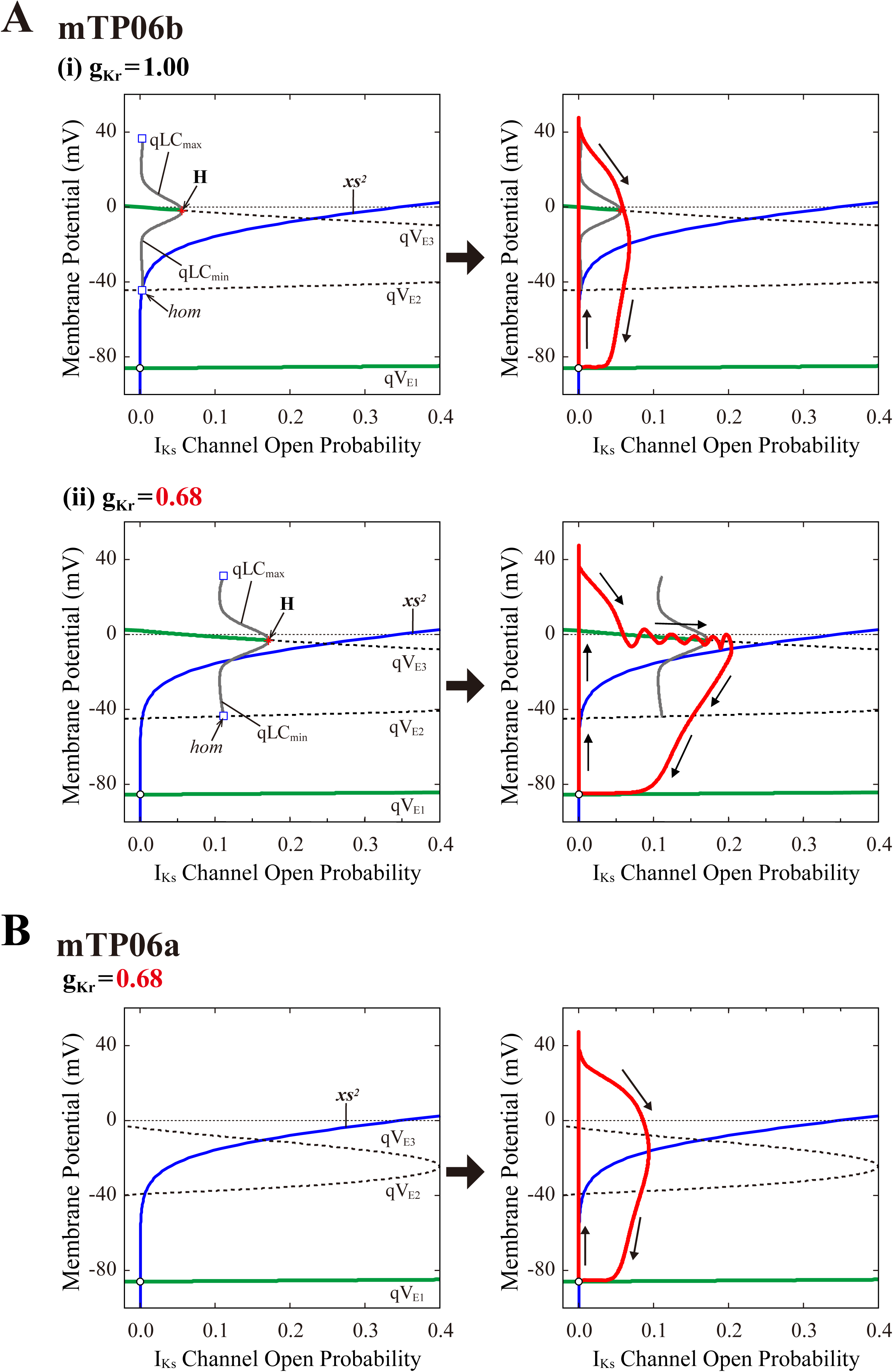
Dynamical mechanisms of EAD initiation and termination determined by the slow-fast decomposition analysis for the mTP06 models. Shown are one-parameter bifurcation diagrams of quasi-equiribrium points (qEPs) and quasi-limit cycles (qLCs), where the steady-state branches as loci of V_m_ at qEPs (qV_E1-3_) and periodic branches as the potential minimum (qLC_min_) and maximum (qLC_max_) of qLCs are depicted as functions of the square of the I_Ks_ activation gating variable (***xs***^2^), i.e. I_Ks_ channel open probability for the fast subsystems of the g_Kr_-normal (**A-(i)**, *left*) and g_Kr_-reduced (**A-(ii)**, *left*) mTP06b model and g_Kr_-reduced mTP06a model (**B**, *left*). Other slow variables, Na_i_ and Ca_SR_, were fixed at constant values: Na_i_ = 6 mM for all cases; Ca_SR_ was fixed at the values which were reached just before occurrence of the first EAD, i.e., at 0.5 mM and 1.5 mM for the normal and g_Kr_-reduced mTP06b model, respectively, and at 0.5 mM for the g_Kr_-reduced mTP06a model. The steady-state branches consist of the stable (green solid lines) and unstable (black dashed lines) segments. The periodic branches (gray solid lines) are all unstable. The blue lines indicate the steady-state ***xs***^2^ curve. Trajectories of the full system (with the fixed Ca_SR_ and Na_i_) are superimposed on the bifurcation diagrams for the fast subsystems (red lines in each right panel). **H**, Hopf bifurcation; ***hom***, homoclinic bifurcation.

#### Dynamical mechanisms of spontaneous SR Ca^2+^ release-mediated EAD

To clarify the dynamical mechanisms of spontaneous SR Ca^2+^ release-mediated EAD formation, we further examined the stability, dynamics and bifurcations of the voltage-clamped mTP06 model. Ca^2+^ dynamics during a train of 1-s depolarizing test pulses to −10 mV (from the holding potential of −85 mV) applied at 2-s intervals to mimic APs evoked by 0.5 Hz pacing were first determined for the mTP06b model with different P_up_ (**Fig. 9A**). Spontaneous SR Ca^2+^ releases occurred when Ca_SR_ increased at higher P_up_, as indicated by the dots in **Fig. 9A**; as P_up_ increased, the time to the first Ca^2+^ release and period of spontaneous Ca^2+^ releases shortened, and their frequency increased. **Figure 9B** shows one-parameter bifurcation diagrams of the steady-state stability and dynamics of Ca_i_ as functions of the clamped-V_m_ in the voltage-clamped mTP06b model. Steady-state intracellular Ca^2+^ concentrations (EPs) in the voltage-clamped model cell were stable at hyperpolarized and depolarized V_m_ (green traces in the right and *middle* panels of **Fig. 9B**) but became unstable via supercritical HBs in the V_m_ range of AP phase 2 and early phase 3 (dashed traces in **Fig. 9B**, middle). LCs emerging from the HB points were first stable but were destabilized via NSBs after small changes in V_m_; spontaneous Ca^2+^ oscillations occurred in the V_m_ range of unstable LCs, i.e., between *NS_1_* and *NS_2_* (gray traces labeled as LC_min_ and LC_max_ in **Fig. 9B**). As shown in **Fig. 9C**, the unstable V_m_ region (*uEP*) was enlarged by increasing P_up_ (see **Fig. 9C**, *left*), decreasing I_NCX_ activity (**Fig. 9C**, *middle*), and/or enhancing I_CaL_ (**Fig. 9C**, *right*), all of which led to increases in Ca_SR_.

**Figure 9:**
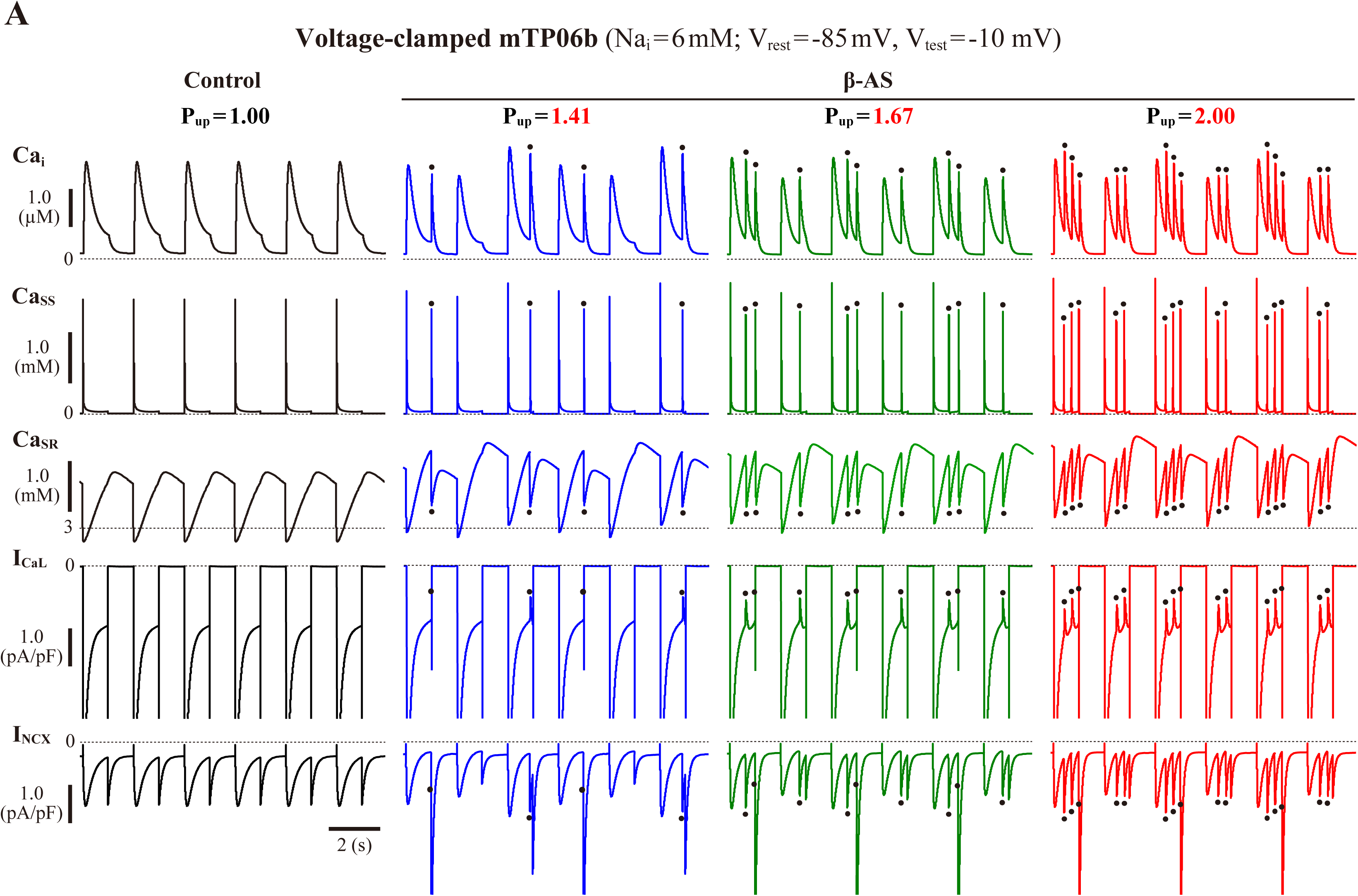
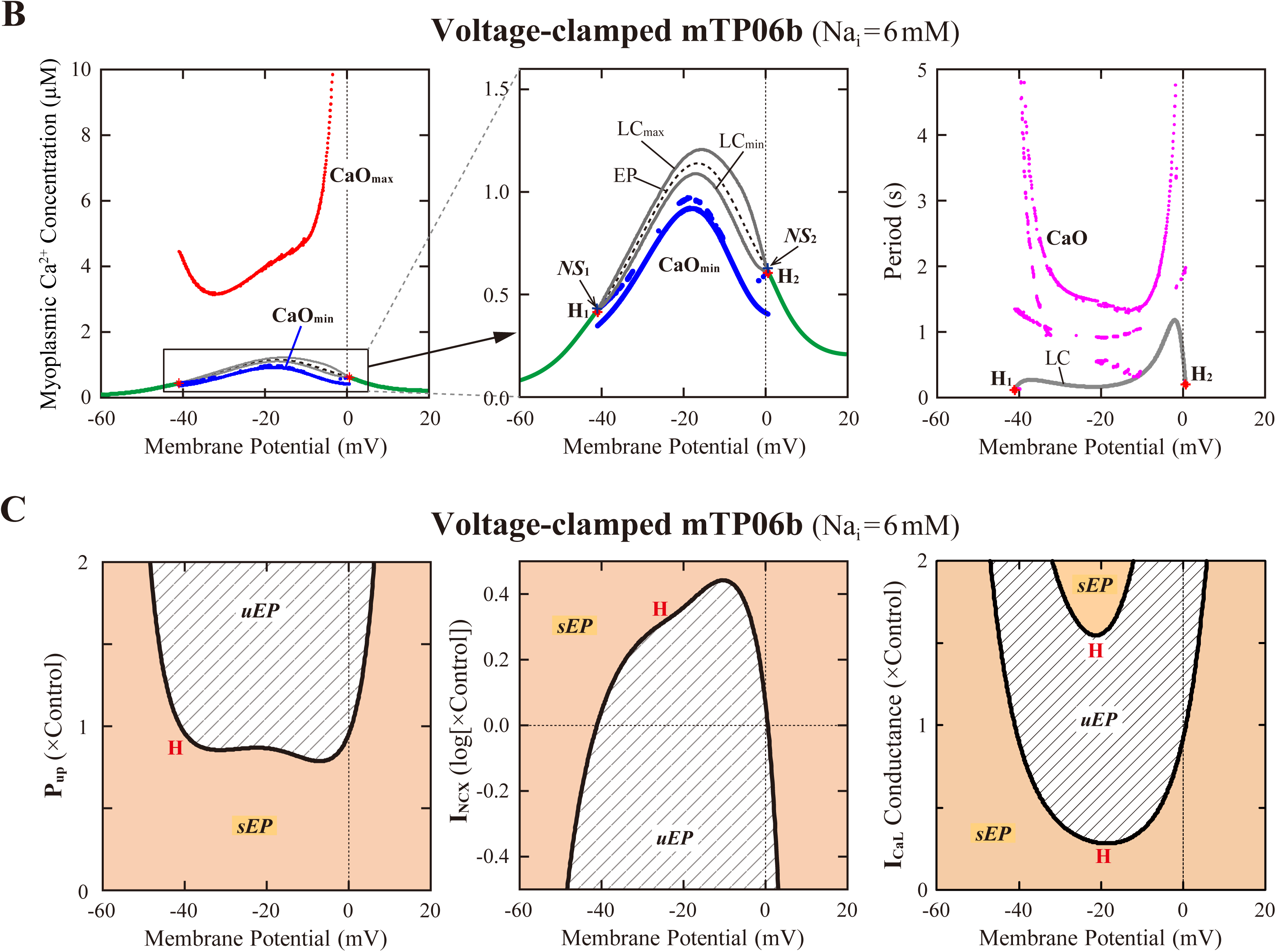
Stability and bifurcations of intracellular Ca^2+^ dynamics in voltage-clamped mTP06b model cell. **(A)** Dynamics of intracellular Ca^2+^ concentrations as well as Ca^2+^-dependent sarcolemmal currents in the Na_i_-fixed voltage-clamped model cell under the control condition (P_up_ = 1) and the conditions of β-AS (P_up_ = 1.41, 1.67 and 2.00). Temporal behaviors of the voltage-clamped model cell were computed for 10 min; Ca_i_, Ca_ss_, Ca_SR_, I_CaL_ and I_NCX_ for the last 12 s (6 pulses) are shown as steady-state dynamics. Spontaneous SR Ca^2+^ releases, as evidenced by abrupt falls of Ca_SR_ and increases in Ca_i_, Ca_ss_ and inward I_NCX_ with attenuated I_CaL_, occurred at higher P_up_ (indicated by the dots). **(B)** One-parameter bifurcation diagrams of the equilibrium point (EP) and extrema of limit cycles (LC_min/max_) and spontaneous Ca_i_ oscillations (CaO_min/max_) as functions of V_m_ for the voltage-clamped model cell (*left* and *middle*). The middle panel shows an enlarged diagram of the rectangular area in the left panel. The periods of spontaneous Ca^2+^ oscillations (CaO) and limit cycles (LC) are also plotted against V_m_ (*right*). **H**_**1–2**_, Hopf bifurcations of the EP; ***NS***_**1–2**_, Neimark-Sacker bifurcations of the LC. **(C)** Two-parameter diagrams on the V_m_–P_up_ (*left*), V_m_–I_NCX_ (*middle*) and V_m_–g_CaL_ (*right*) planes, indicating how the unstable V_m_ range changed depending on P_up_, I_NCX_ and I_CaL_. HB values, i.e., the critical V_m_ at which an EP is (de)stabilized are plotted as functions of P_up_, I_NCX_ and I_CaL_ for the Na_i_-fixed mTP06b model; the HB points were very close to the NSB points at which LCs were destabilized with the emergence of CaOs.

To further clarify the Ca_SR_-dependent mechanism of spontaneous SR Ca^2+^ releases in the P_up_-increased mTP06b model and why SR Ca^2+^ release-mediated EADs emerge more frequently at larger P_up_, we also performed the slow-fast decomposition analysis for the slow variable Ca_SR_. Bifurcation diagrams were constructed as functions of Ca_SR_ for the voltage-clamped fast subsystem composed of the voltage-independent state variables f_CaL_ (Ca^2+^-dependent inactivation gate for I_CaL_), R (proportion of closed SR Ca^2+^ release channels), Ca_ss_, and Ca_i_ (**Fig. 10**). Trajectories of the voltage-clamped full system dynamics as shown in **Fig. 9A** for the normal (1) and enhanced (1.67) P_up_ were superimposed on the diagrams. The steady states of the fast subsystem, stable at lower Ca_SR_ (green traces in the middle and right panels of **Fig. 10**), became unstable via an HB at higher Ca_SR_ (dashed traces in the middle and right panels of **Fig. 10**). In the P_up_-enhanced system, spontaneous SR Ca^2+^ releases as shown in blue trajectories in the middle and right panels of **Fig. 10B** occurred when the full system trajectory, moving along the stable steady-state branch, passed through the HB point, i.e., when Ca_SR_ exceeded the HB value. In contrast, the P_up_-normal system did not exhibit spontaneous SR Ca^2+^ release, because an increment of Ca_SR_ (Ca^2+^ refilling of the SR) during Ca^2+^ transient decay was too slow for the full system trajectory to reach the HB point for Ca_SR_ before V_m_ repolarization (**Fig. 10A**, *middle* and *right*).

**Figure 10:**
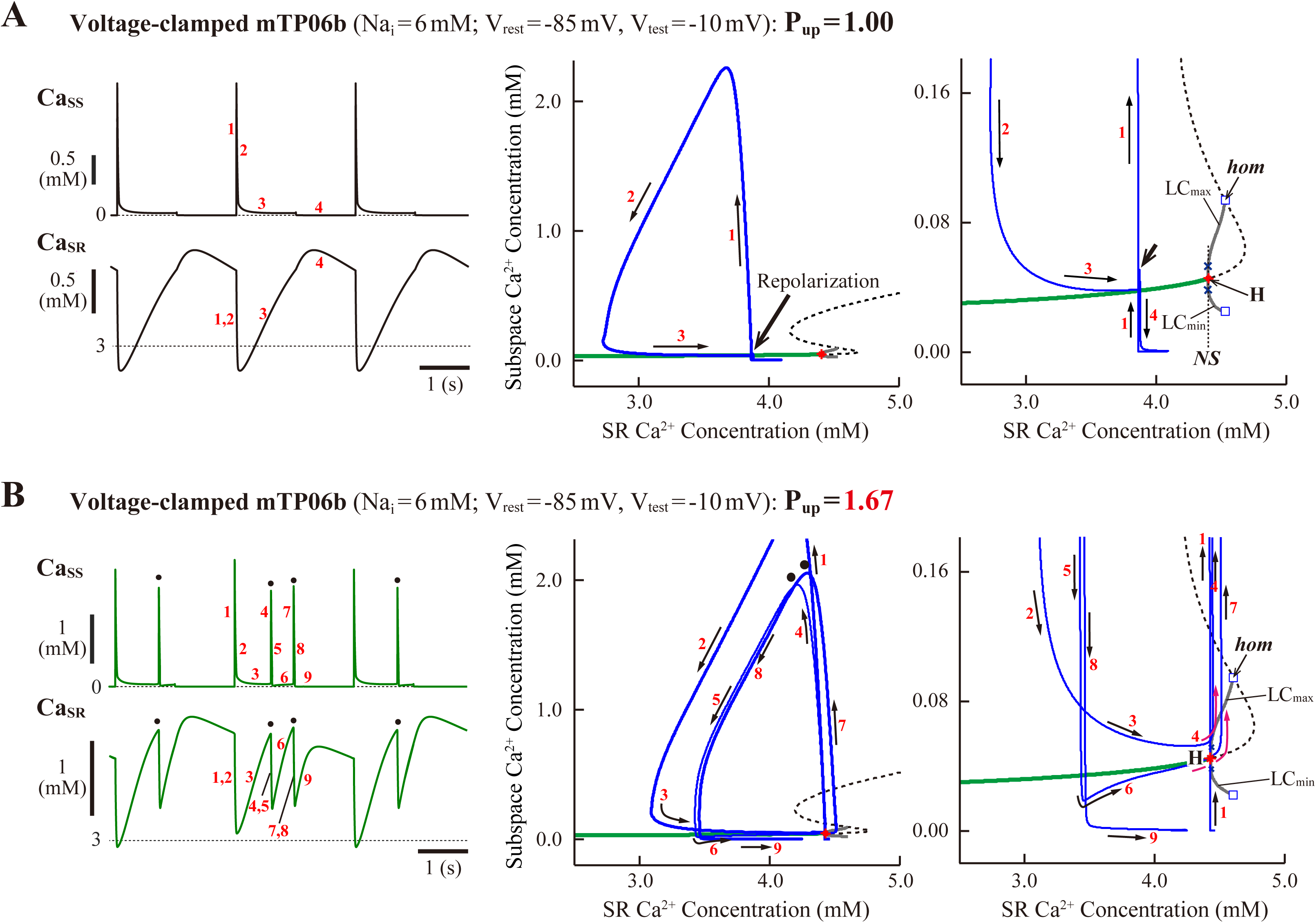
Slow-fast decomposition analysis for the slow variable Ca_SR_ of the voltage-clamped mTP06b model. Normal (*middle*) and expanded (*right*) scale views of one-parameter bifurcation diagrams, where the steady-state branch as a locus of Ca_ss_ at EPs, as well as periodic branches as the minimum (LC_min_) and maximum (LC_max_) of Ca_ss_ limit cycles (LCs) that were almost always unstable, are plotted as functions of Ca_SR_ for the fast subsystems of the voltage-clamped model cells with the normal **(A)** and increased **(B)** P_up_. The green solid and black dashed lines represent stable and unstable steady-state branch, respectively. Subcritical HB (**H**) with the emergence of an unstable LC, NSB from a stable LC to an unstable one following a SNB of LCs (***NS***), and homoclinic bifurcation (***hom***) are located. Trajectories projected onto the Ca_SR_-Ca_ss_ plane of the full system during a 1-s step depolarization to −10 mV (from −85 mV) and subsequent 1-s repolarization to −85 mV are superimposed on the diagrams; Ca_ss_ and Ca_SR_ dynamics in the full system during this voltage-clamp pulse are shown on the left. The labels “1”–“9” for the arrows on each diagram (*middle*, *right*) indicate the time course of changes in the state variables (Ca_ss_ and Ca_SR_) along the trajectories, with the labels also given for the Ca_ss_ and Ca_SR_ dynamics (*left*).

## Discussion

In this study, we theoretically investigated dynamical mechanisms of EAD formation in the TP06 model for HVMs, which has often been used for simulations and theoretical analyses of reentrant arrhythmias, automaticity, multi-stability and EAD formation in HVMs, in relation to the model cell dynamical behaviors and their bifurcations. Our findings are summarized as follows: 1) The M cell version of the TP06 model with accelerated I_CaL_ inactivation (mTP06b model) could reproduce EADs in the I_Ks_-reduced LQT1-type and I_Kr_-reduced LQT2-type HVMs, including β-AS-induced EADs in LQT1 and bradycardia-dependent EADs in LQT2 (**Figs. 2–5** for validation of the model); 2) two types of EADs with distinct initiation mechanisms, i.e., I_CaL_ reactivation–dependent EADs (**Figs. 1**, **2A-(a)** and **6B**) and spontaneous SR Ca^2+^ release-mediated EADs (**Figs. 2A-(b)**, **5A** and **6B**) were detected, the latter not reproduced by the K05 or O11 model; 3) SR Ca^2+^ uptake/release machinery (intracellular Ca^2+^ dynamics), I_NCX_, and I_CaL_ strongly affected EAD generation via modulating stability of the model cell dynamical behaviors and their bifurcations in a manner different from that in the K05 or O11 model (**Figs. 6** and **7**); 4) during pacing, EAD termination prior to the next stimulus was I_Ks_ activation–dependent, I_Ks_ being indispensable for AP repolarization (**Figs. 1**, **2**, **3** and **8**); 5) spontaneous SR Ca^2+^ releases and Ca^2+^ oscillations occurred with their incidence and frequency increased by enhancing SR Ca^2+^ uptake, attributable to the instability of intracellular Ca^2+^ dynamics as unveiled by voltage clamping (**Figs. 9** and **10**). In summary, EAD formation and its dynamics in the paced (non-autonomous) mTP06 model cell basically depended on stability and bifurcations of the non-paced (autonomous) model cell. Bifurcation phenomena and dynamical mechanisms of EAD formation in the mTP06 model were different from those in the K05 and O11 models tested previously (Kurata *et al*. 2017) in several respects.

### Validation of the mTP06 Model for EAD Reproducibility in LQT1 and LQT2 Conditions

We investigated whether and how the modified TP06 models could recapitulate dynamic properties of LQT1 and LQT2 HVMs to cause EADs. The mTP06a and mTP06b models were first tested for EAD formation associated with bifurcation phenomena during inhibition of I_Kr_ and/or I_Ks_ and enhancement of I_CaL_ (β-AS) by constructing bifurcation diagrams, as well as by numerical simulations (**Figs. 2–5**).

### EAD formation in LQT1 and LQT2 conditions (mTP06a vs. mTP06b)

I_Ks_ and I_Kr_ play pivotal roles in preventing EADs, with their reduction causing LQT1 and LQT2, respectively (Shimizu & Horie, 2011; Shimizu, 2013). Like the K05 and O11 models, the modified TP06 models could recapitulate EAD formation in the I_Ks_-reduced LQT1-type and I_Kr_-reduced LQT2-type HVMs. The mTP06b model with accelerated I_CaL_ inactivation was much more vulnerable to EAD formation than the mTP06a model, consistent with the previous experimental finding that slowing I_CaL_ inactivation eliminated EADs (Qu *et al*. 2013). As demonstrated by the slow-fast decomposition analysis (**Fig. 8**), higher susceptibility of the mTP06b model is attributable to the stabilization of qEPs at depolarized V_m_ close to the plateau V_m_ in the ***xs***-parameterized fast subsystem by accelerating I_CaL_ inactivation. Modulating I_Ks_ shifted the critical points of EAD formation (red thick solid line in **Fig. 3A**) and HB (black thin solid line in **Fig. 3A**) during I_Kr_ inhibition in a manner very similar to those in the K05 and O11 models (Kurata *et al*. 2017).

### Rate dependence of EAD formation (validation for LQT2 model)

In LQT2 patients, fatal cardiac events often occur during sleep or at rest, i.e., when I_Ks_ activation would delay in bradycardia (Shimizu & Horie, 2011; Shimizu, 2013). The K05 and O11 models could partially reproduce the bradycardia-related EADs (Kurata *et al*. 2017). To clarify whether the mTP06b model could recapitulate bradycardia-related EAD formation in LQT2-type HVMs, we tested the pacing rate dependence of EAD formation during I_Kr_ reduction in the mTP06b model. The Na_i_-variable mTP06b model could partly reproduce the rate-dependent EAD generation in LQT2 patients (**Fig. 4**). In our previous study for the Na_i_-variable K05 and O11 models (Kurata *et al*. 2017), the facilitation of EAD formation during lower rate pacing was accompanied by the decrease in Na_i_ and resulting reductions in outward I_NaK_ and inward shift of I_NCX_. This study have also demonstrated for the mTP06b model that the facilitation of EAD formation at lower pacing rates is mainly due to the decrease in Na_i_ and resultant changes in I_NaK_ (**Fig. 4B**). Thus, the major mechanism for bradycardia-related EADs in the mTP06b model is essentially the same as those in the K05 and O11 models.

### EAD formation during β-AS (validation for LQT1 model)

In LQT1 patients with smaller I_Ks_, fatal cardiac events are exercise-induced (tachycardia-related), for adrenergic enhancement of I_CaL_ is no longer counterbalanced by the concomitant stimulation of I_Ks_; the smaller increase in I_Ks_ leads to the occurrence of EADs that trigger ventricular tachyarrhythmia. The K05 model, but not the O11 model, could reproduce this I_Ks_ reduction-related EAD formation as a cause of ventricular tachycardia in LQT1 patients during β-AS (Kurata *et al*. 2017). Thus, we further tested whether the mTP06a/b models could recapitulate facilitated EAD formation under the conditions of β-AS in LQT1-type HVMs. In the mTP06b model with normal I_Ks_, EAD formation during I_CaL_ enhancement via β-AS was avoided by the concomitant increase in I_Ks_; I_Ks_-reduced LQT1-type HVMs exhibited EADs with relatively small enhancement of I_CaL_ via falling into the area of EAD generation (RF and fR regions in **Fig. 5B**). Thus, the mTP06b model could also reproduce β-AS-related EAD formation in LQT1 cardiomyocytes, which could clearly be accounted for by the I_CaL_- and I_Ks_-dependent bifurcation properties of the model cell. However, EAD formation in the mTP06b model during β-AS was different in a mechanism from that in the K05 model: EADs in the mTP06b model involved spontaneous SR Ca^2+^ releases and resulting increments of inward I_NCX_ (**Fig. 5A** and **Online Supplementary Fig. S4**), while those in the K05 model solely I_CaL_-reactivation dependent, not involving spontaneous SR Ca^2+^ releases (Kurata *et al*. 2017). More sophisticated HVM models incorporating β-AS-related modulating factors, like those developed by Saucerman et al. (2003) and Kuzumoto et al (2008), are required for further investigations of the mechanisms of exercise-induced EAD formation in LQT1 HVMs.

### Dynamical Mechanisms of Phase-2 EAD Formation in the mTP06 Models

#### Initiation mechanisms

At least two mechanisms appeared to underlie the initiation of phase-2 EADs in the mTP06b model: 1) I_CaL_ reactivation-dependent mechanism which operates and causes EADs even in the absence of spontaneous SR Ca^2+^ releases at lower P_up_ and lower pacing rates, and 2) spontaneous SR Ca^2+^ release-mediated mechanism activating inward I_NCX_ at higher P_up_ and higher pacing rates (**Figs. 2A-(b)** and **6B**).

Qu *et al*. (Qu & Chung, 2012; Qu *et al*. 2013) suggested major contribution of I_CaL_, which plays a pivotal role in creation of an unstable EP (qEP) at plateau V_m_ by its window current and gating kinetics. This mechanism could also be applicable to the HVM models, as discussed in our previous article (Kurata *et al*. 2017). The major contribution of I_CaL_ to EAD formation was also suggested in many previous experimental and theoretical studies for ventricular myocytes (Guo *et al*. 2007; Yamada *et al*. 2008; Xie *et al*. 2010; Corrias *et al*. 2011; Madhvani *et al*. 2011; Chang, Chang *et al*. 2012; Chang, Sato *et al*. 2012; Milberg *et al*. 2012; Qu & Chung, 2012). EAD formation in the mTP06 model with lower P_up_ is also attributable to I_CaL_ reactivation in that reactivated I_CaL_ contributes to V_m_ depolarization (**Figs. 1**, **2A-(a)**, **6B**, and **Online Supplementary Fig. S5**). The slow-fast decomposition analysis of the guinea-pig ventricular myocyte model have suggested that the I_CaL_-dependent destabilization of a qEP and formation of a stable qLC via an HB in the fast subsystem is required for EAD generation in the full system (Tran *et al*. 2009; Qu *et al*. 2013; Song *et al*. 2015). In the mTP06b model, however, transient trapping of the full system trajectory occurred around the stable and unstable qEPs without forming a stable qLC, indicating that the emergence of a stable qLC is not necessarily needed for EAD formation. As mentioned above, the initiation of EADs in the mTP06b model is attributable to the stabilization of qEPs at depolarized V_m_ in the ***xs***-parameterized fast subsystem, which causes transient trapping of the full system trajectory around the stable qEP; I_Kr_ reduction promotes EAD formation by broadening the region of stable qEPs at depolarized V_m_ (**Fig. 8**).

As another possible mechanism for EAD initiation, many recent experimental studies have strongly suggested the spontaneous SR Ca^2+^ release causing Ca_i_ oscillations, oscillatory increases in inward I_NCX_, and resulting V_m_ depolarization during β-AS (Choi *et al*. 2002; Volders *et al*. 2003; Zhao *et al*. 2012) and in I_Kr_-reduced LQT2 cardiomyocytes (Kim *et al*. 2005; Němec *et al*. 2010, 2016), which is similar to the mechanism for delayed afterdepolarizations (DADs) induced by spontaneous SR Ca^2+^ releases under Ca^2+^ overload conditions or β-AS (e.g., Volders *et al*. 2003; Zhao *et al*. 2012) and the Ca^2+^ clock mechanism for sinoatrial node cell pacemaking (Maltsev & Lakatta, 2009). The K05 or O11 model could not reproduce the spontaneous SR Ca^2+^ release as a cause of phase-2 EADs (Kurata *et al*. 2017). In contrast, the mTP06b model could clearly replicate this scenario in a Ca_SR_-dependent manner (**Figs. 2A-(b)**, **5A**, **6B**, and **10**), while it was not found in the previous study using a modified TP06 model (Vandersickel *et al*. 2014). Such SR Ca^2+^ release-mediated EADs under β-AS conditions (**Fig. 5**) have also been reproduced by the rabbit ventricular myocyte model (Volders *et al*. 2000; Song *et al*. 2015). In addition, very similar formulas for SR Ca^2+^ handling in the Grandi et al model (Grandi *et al*. 2010) suggest that spontaneous SR Ca^2+^ releases to cause EADs also occur in this HVM model, as suggested previously (Trenor *et al*. 2013).

This study further suggests that the occurrence of spontaneous SR Ca^2+^ releases and Ca^2+^ oscillations are attributable to instability of intracellular Ca^2+^ concentrations in a steady state, destabilization of which leads to spontaneous Ca^2+^ oscillations (**Figs. 9** and **10**). This scenario, i.e., steady-state destabilization for spontaneous Ca^2+^ oscillations involving ryanodine or IP_3_ receptors, has previously been suggested by bifurcation analyses for cardiac myocytes (Keizer & Levine, 1996; Tveito *et al*. 2012) and for other cells (Schuster *et al*. 2002; Higgins *et al*. 2006; Kusters *et al*. 2007). However, the Ca^2+^ oscillations reported in these previous studies were much longer in period than those observed in the mTP06b model, not relating to EAD formation. To the best of our knowledge, this is the first report demonstrating instability of steady-state intracellular Ca^2+^ concentrations and resulting spontaneous SR Ca^2+^ releases that cause EADs in the HVM model. Himeno et al (Himeno *et al*. 2013) recently developed a novel HVM model incorporating more detailed descriptions of SR Ca^2+^ release mechanisms. This model may be useful for providing more profound insights into how EADs are initiated via spontaneous SR Ca^2+^ releases under certain conditions.

Other candidates to initiate EADs include the reactivation of I_Na_ (Edwards *et al*. 2014); however, I_Na_ was not involved in EAD formation in the HVM models tested.

#### Termination mechanisms

With respect to the mechanisms of the termination of phase-2 EADs (AP repolarization), EADs in the mTP06b model during pacing appeared to terminate in an I_Ks_ activation-dependent (or stimulus-dependent) manner: The open probability of I_Ks_ channels (***xs***^2^) increased progressively in the model cell with relatively small I_Kr_, i.e., LQT2-like cells (**Fig. 2A-(a)**). Tran *et al*. (2009) suggested the major role of the slow I_Ks_ activation for the guinea-pig ventricular myocyte model by the slow-fast decomposition analysis in which the slow I_Ks_ activation gating variable was assumed to be a parameter for the fast subsystem. In the diagram for the slow gating variable-parameterized fast subsystem with a superimposed full system trajectory, gradual increases in the slow variable led the full system trajectory slowly across the stable steady-state branch of qEP and then into the region of the stable periodic branch of qLC through an HB point, resulting in the termination of EADs via a homoclinic bifurcation of qLC (Tran *et al*. 2009; Qu *et al*. 2013; Song *et al*. 2015). This scenario, known as the Hopf-homoclinic bifurcation mechanism, appears to be applicable to EAD formation in the K05 model for HVMs as well (Kurata *et al*. 2017). Consistent with these previous reports, the mTP06b model exhibited slow I_Ks_ activation-dependent EADs in the ***xs***^**2**^ regions of stable and unstable qEPs (**Fig. 8A**); however, a stable qLC region or homoclinic bifurcation to yield EAD termination was not detected for the fast subsystem of the mTP06b model. EAD terminated simply via the destabilization of a qEP in the I_Kr_-reduced mTP06b model, suggesting that the Hopf-homoclinic bifurcation scenario is not necessarily applicable. When I_Kr_ was smaller, EADs did not terminate but AP repolarization failure (arrest at depolarized V_m_) occurred, because the full system trajectory reached equilibrium at the intersection of the stable steady-state branch (qV_E3_) and steady-state ***xs***^**2**^ curve.

In the I_Ks_-eliminated system, EADs would not terminate unless there exist other slow components or factors, such as the slowly-inactivating I_CaL_ or late I_Na_ and intracellular Na^+^ accumulation to increase outward I_NaK_ gently. Our previous study using the K05 and O11 models indicated that EAD termination might occur in a slow I_CaL_ inactivation-dependent manner when I_Ks_ was relatively small (Fig. 3 in Kurata *et al*. 2017). However, I_CaL_ inactivation-dependent EAD termination was not clearly detected in the TP06 model. Other candidates for slow variables to cause EAD termination include the slow inactivation of late I_Na_ (Horvath *et al*. 2013; Trenor *et al*. 2013; Asakura *et al*. 2014) not incorporated into the TP06 model and gradual increases in Na_i_ (Chang *et al*. 2012; Xie *et al*. 2015). After cessation of pacing, the I_Kr_-reduced and/or I_CaL_-enhanced TP06b model could exhibit long-term EAD bursts the termination of which was induced by slow elevation of Na_i_ and resulting enhancement of outward I_NaK_ (data not shown). This Na_i_-dependent mechanism has previously been demonstrated for a rabbit ventricular AP model as well (Chang *et al*. 2012). Nevertheless, the slow Na_i_ elevation (intracellular Na^+^ accumulation) is unlikely as a termination mechanism for short-term EADs during pacing at 0.2–2 Hz in the TP06 model.

### Comparisons with Other HVM Models for Bifurcation Phenomena and EAD Mechanisms

#### Roles of I_Kr_ and I_Ks_

I_Ks_/I_Kr_-dependent bifurcations and EAD formation of the mTP06b model occurred in essentially the same way as those in the K05 and O11 models (Kurata *et al*. 2017). On the g_Ks_–g_Kr_ parameter planes, decreasing I_Ks_ shifted the critical points of EAD emergence to promote EAD formation during inhibition of I_Kr_, and vice versa, as in the other HVM models (Kurata *et al*. 2017). However, the I_Ks_/I_Kr_-dependent properties of the mTP06 model (**Figs. 2** and **3**) were different in the following respects: 1) on the g_Ks_–g_Kr_ parameter plane, the shifts in HB points depended more strongly on g_Ks_ than on g_Kr_, while more strongly on g_Kr_ in the O11 model; and 2) a stable LC (periodic branch) as observed in the parameter region of EAD formation in the K05 and O11 models (Fig. 2 in Kurata *et al*. 2017) was not detectable. In addition, one of the major differences is that the mTP06 model requires I_Ks_ for EAD termination, i.e., repolarization failure occurred abruptly during I_Kr_ inhibition or I_CaL_ enhancement when I_Ks_ was absent or small, whereas I_Ks_ was not necessarily needed in the K05 or O11 model; I_CaL_ inactivation-dependent EAD termination was possible in these HVM models, but not in the mTP06 model.

#### Role of I_CaL_

It is well known that reactivation of I_CaL_ contributes to the generation of phase-2 EADs (January & Riddle, 1989; Ming *et al*. 1994). Previous theoretical studies using the guinea-pig and human ventricular myocyte models (Tran *et al*. 2009; Qu *et al*. 2013; Kurata *et al*. 2017) suggest that I_CaL_ contributes to EAD formation via the inductions of a HB to destabilize an EP (qEP) and a stable LC (qLC), and that accelerating I_CaL_ inactivation or slowing I_CaL_ activation prevents the HB-induced destabilization of EP (qEP), broadens the g_CaL_ and g_Kr_ regions of stable V_E3_ (qV_E3_), shrinks LC (qLC) oscillations, and thereby prevents the emergence of EADs. In our previous study using the K05 and O11 models (Kurata *et al*. 2017), as g_CaL_ increased, the non-paced autonomous model cell systems underwent the emergence of stable LCs with the g_CaL_ regions of EADs overlapping the regions of the stable LC. EAD formation in the mTP06 model at lower P_up_ is also I_CaL_-dependent, for I_CaL_ contributes to V_m_ upstroke in EADs. However, the ways of I_CaL_-dependent bifurcations and EAD formation in the mTP06 model were different from those in the other HVM models in that a stable LC, the parameter region of which overlapped that of EADs in the K05 and O11 models (**Fig. 4** in Kurata *et al*. 2017), was not detectable in the mTP06 model (**Fig. 7**). In the K05 and O11 models, EADs often emerged in the vicinity of the critical point at which a stable LC appeared during I_CaL_ increases (Kurata *et al*. 2017), suggesting that EAD formation depends on I_CaL_ responsible for the instability of EPs and generation of stable LCs. EADs also occurred in the I_CaL_-enhanced mTP06b model when unstable LCs emerged, but stable LCs were not detected (**Fig. 7A**). I_CaL_/I_Ks_-dependent properties of EAD formation in the mTP06a and mTP06b models are very similar to those in the O11 and K05 model, respectively, while I_CaL_/I_Ks_-dependent HB properties of the mTP06 models were totally different (**Fig. 7B** and **Online Supplementary Fig. S8**). The mTP06b model as well as the K05 model, but not the mTP06a or O11 model, was capable of reproducing I_CaL_-dependent EAD formation during β-AS in LQT1 (**Fig. 5**). In the mTP06b model, I_CaL_ further contributed to spontaneous SR Ca^2+^ releases via the enhancement of the instability of intracellular Ca^2+^ dynamics (**Fig. 9C**).

#### Roles of SR Ca^2+^ uptake/release and I_NCX_

The effects on EAD formation of modulating SR Ca^2+^ uptake/release and I_NCX_ in the mTP06 model were different from those in the other HVM models tested previously (Kurata *et al*. 2017). With respect to the onset of EADs during I_Kr_ inhibition or I_CaL_ enhancement, our previous study using the K05 model showed that enhancing SR Ca^2+^ uptake/release facilitated EAD generation via an inward shift of I_NCX_ and reduction of I_CaL_ which lowered the plateau V_m_ and thereby limited I_Ks_ activation during the preconditioning phase before EAD formation (Kurata et al. 2017). In the mTP06 model, however, the effects of modulating SR Ca^2+^ uptake/release (changing P_up_ from the control value) on the onset of EADs were opposite to those in the K05 model, while similar to those in the O11 model; inhibition of SR Ca^2+^ uptake/release or Ca^2+^ transient did not prevent but slightly facilitated EAD formation during I_Kr_ inhibition or I_CaL_ increments (**Fig. 6A**). This inconsistency is mainly due to the differences in SR Ca^2+^ uptake/release mechanisms and Ca_i_-dependent behaviors of I_CaL_ and I_NCX_. In regard to the effect on EP stability, the three HVM models are completely different: SR Ca^2+^ cycling destabilized EPs and induced spontaneous oscillations in the mTP06 model (**Fig. 6** and **Online Supplementary Fig. S6**), exerted no effect on EP stability (and little affected LC oscillation) in the K05 model, and broadened the parameter regions of stable EPs in the O11 model. Thus, the influences of SR Ca^2+^ cycling on bifurcations and EAD formation are model dependent; further studies using more sophisticated HVM models are required.

One of the prominent properties of the mTP06 model is the instability of steady-state intracellular Ca^2+^ concentrations resulting in the spontaneous SR Ca^2+^ release as a cause of phase-2 EADs that occur at higher P_up_ to increase Ca_SR_ (**Figs. 9** and **10**). Spontaneous SR Ca^2+^ releases and Ca^2+^ oscillations were suggested to induce EADs via enhancing inward I_NCX_ under certain conditions such as β-AS (Volders *et al*. 2000; Zhao *et al*. 2012) and I_Kr_ inhibition in LQT2 (Choi *et al*. 2002; Kim *et al*. 2005; Němec *et al*. 2010, 2016). However, the K05 or O11 model did not exhibit spontaneous SR Ca^2+^ releases even at higher P_up_, because steady-state intracellular Ca^2+^ concentrations were always stable independently of V_m_; although Ca_i_ oscillations occurred during EADs in the K05 and O11 models, these Ca_i_ oscillations were not induced by spontaneous SR Ca^2+^ releases but by oscillatory reactivation of I_CaL_ (Kurata *et al*. 2017). Thus, the mTP06 model but not the other HVM models tested previously could reproduce the spontaneous SR Ca^2+^ release as a cause of phase-2 EADs.

In previous experimental studies, enhancing I_NCX_ tended to promote EADs (Pott *et al*. 2012), while inhibition of I_NCX_ prevented EADs (Nagy *et al*. 2004; Milberg *et al*. 2012). The mTP06 model predicted facilitation of EAD formation by enhanced I_NCX_, consistent with the experimental finding (Pott *et al*. 2012). This promotion of EAD generation was not only via enhancing inward I_NCX_ but also via secondary reductions in I_CaL_ inactivation and resultant increases of I_CaL_ window currents, as in the K05 and O11 models (Kurata *et al*. 2017). In the mTP06 model, the suppression of phase-2 EADs was yielded only by small inhibition of I_NCX_, with greater inhibition of I_NCX_ leading to facilitated EAD formation; as in the other HVM models, small inhibition of I_NCX_ prevented EADs via primary reductions of inward I_NCX_ and secondary decreases of I_CaL_.

#### Limitations and Perspectives of Study

In this study, we employed bifurcation analyses to investigate the dynamical mechanisms of EAD generation in a HVM model, finding how individual ionic channel and transporter currents, as well as intracellular Ca^2+^ dynamics, contribute to the initiation, termination and modulation of EADs. As summarized in our preceding article (Kurata *et al*. 2017), bifurcation analyses have been used for elucidating the dynamical mechanisms of sinoatrial node pacemaking, abnormal automaticity in ventricular myocytes, generation of biological pacemaker activity, and EAD formation in ventricular myocytes. These theoretical studies have clearly demonstrated the significance of bifurcation analyses for general understanding and systematic description of the dynamical mechanisms of normal and abnormal oscillatory behaviors.

There are many limitations of our study including incompleteness of the HVM model, as well as the lack of experimental evidence for bifurcation phenomena in real HVMs. More sophisticated HVM models have to be used for more detailed theoretical investigations. Recently developed models incorporating Markovian state schemes for voltage-gated channels, such as Iyer *et al*. (2004), Asakura *et al*. (2014), and Himeno *et al*. (2015) models, may be useful. Moreover, incorporation of more elaborate schemes for the mechanisms of SR Ca^2+^ release and intra-SR Ca^2+^ transfer (Laver, 2007, 2009; Chen *et al*. 2014) would also be crucial. Our preceding (Kurata *et al*. 2017; Tsumoto *et al*. 2017) and present studies have demonstrated that EAD mechanisms are different depending on models and parameter values. As suggested by Zaniboni (Zaniboni *et al*. 2010; Zaniboni, 2012) and demonstrated by the modifications of the TP06 model in this study, different parameter sets for a model cell can yield very similar AP waveforms with different sensitivities to parameter changes; therefore, we have to test as many models as possible for providing more profound understanding of EAD mechanisms.

In this study, bifurcation analysis was limited to a single cell model. However, EAD-related arrhythmias are suggested to be induced by synchronization of EADs in multiple cells (Sato *et al*. 2009; Xie *et al*. 2010); because of electrotonic interactions, EAD formation in multicellular or tissue models may be very different in conditions from that in single cell models (Gibbs *et al*. 1994; Huelsing *et al*. 2000; Weiss *et al*. 2010; Corrias *et al*. 2011). Therefore, we need investigations of the mechanisms for EAD formation and for triggering arrhythmias in human ventricles *in vivo*, which require 1–3 dimensional multicellular (tissue) models, like those used in previous simulation studies (Weiss *et al*. 2010; de Lange *et al*. 2012; Vandersickel *et al*. 2014; Chang *et al*. 2015; Liu *et al*. 2018). Despite many limitations, our studies provide significant insights into the dynamical mechanisms of EAD generation in LQT1 and LQT2 HVMs by utilizing recently developed HVM models.

## Author Contributions

Ya.K. conception and design of research; Ya.K. and K.T. performed programing, simulations and bifurcation analyses; Ya.K. and K.T. analyzed data; Ya.K., K.T., and K.H. interpreted results; Ya.K., K.T., M.T., and Yu.K. prepared figures; Ya.K. drafted manuscript; Ya.K., K.T., K.H., and I.H. edited and revised manuscript; Ya.K., K.T., K.H., I.H., M.T., and Yu.K. approved final version of manuscript.

## Supplementary Figure Legends

**Supplementary Figure S1:**
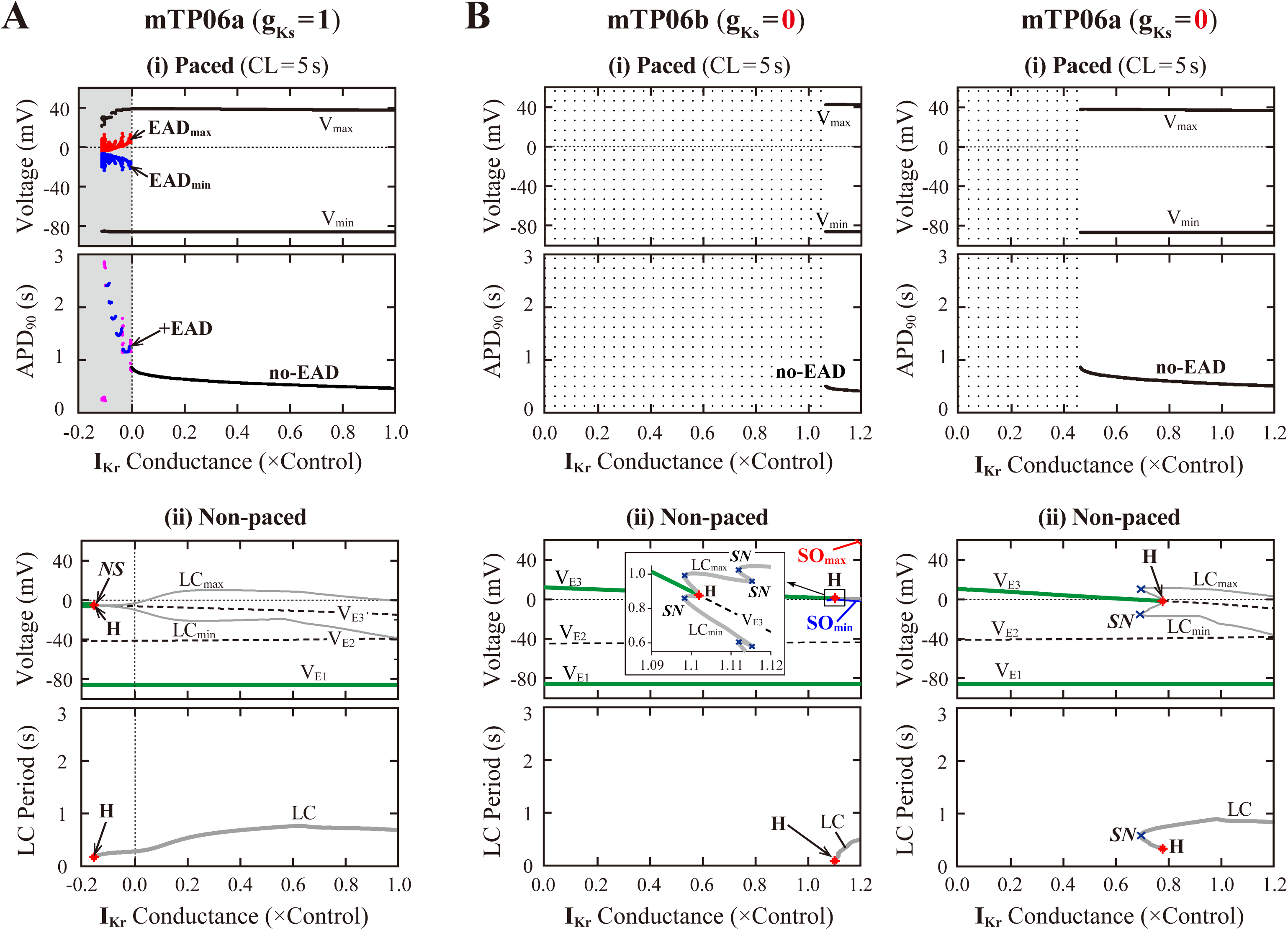
I_Kr_-dependent EAD generations and bifurcations in the I_Ks_-normal and I_Ks_-eliminated mTP06a/b models. Potential extrema of simulated action potential (AP) and EADs, and the AP duration (APD) measured at 90% repolarization (APD_90_) are plotted as functions of normalized g_Kr_ for the I_Ks_-normal mTP06a **(A)** and I_Ks_-removed mTP06a/b **(B)** model cells paced at 0.2 Hz **(i)**. AP dynamics were computed for 1 min at each g_Kr_ value, which was reduced from 1.0 to −0.2 or 1.2 to 0 at an interval of 0.001. The minimum V_m_ during AP phase 4 (V_min_) and the maximum V_m_ during AP phase 2 before EAD formation (V_max_) are represented by black dots for rhythmic APs, and by orange (V_min_) and light green (V_max_) dots for arrhythmic APs. When EADs appeared, their local potential minimum (EAD_min_) and maximum (EAD_max_) were plotted by blue and red dots, respectively. In the diagram for APD_90_, the black, blue and magenta dots represent APD_90_ values for regular APs without EAD, regular APs with EADs, and arrhythmic APs with EADs, respectively (no-EAD: APs without EAD, +EAD: APs with EADs). One-parameter bifurcation diagrams with the steady-state branches as loci of V_m_ at equilibrium points (V_E1-3_) and periodic branches as the potential minimum (LC_min_) and maximum (LC_max_) of limit cycles (LCs) as well as the period of LCs plotted against g_Kr_ are also shown for the non-paced mTP06a/b model cells **(ii)**. The steady-state branches consist of the stable (green solid lines) and unstable (black dashed lines) segments; the periodic branches are almost always unstable, while stable between the Hopf bifurcation (H) and Neimark-Sacker bifurcation (NS) points in Panel A-(ii). The inset in Panel B-(ii) for the I_Ks_-removed mTP06b model is the expanded scale view of the vicinity of the Hopf bifurcation point (rectangular area). Representations and symbols are the same as in Fig. 2B. ***SN***, saddle-node bifurcation of LCs.

**Supplementary Figure S2:**
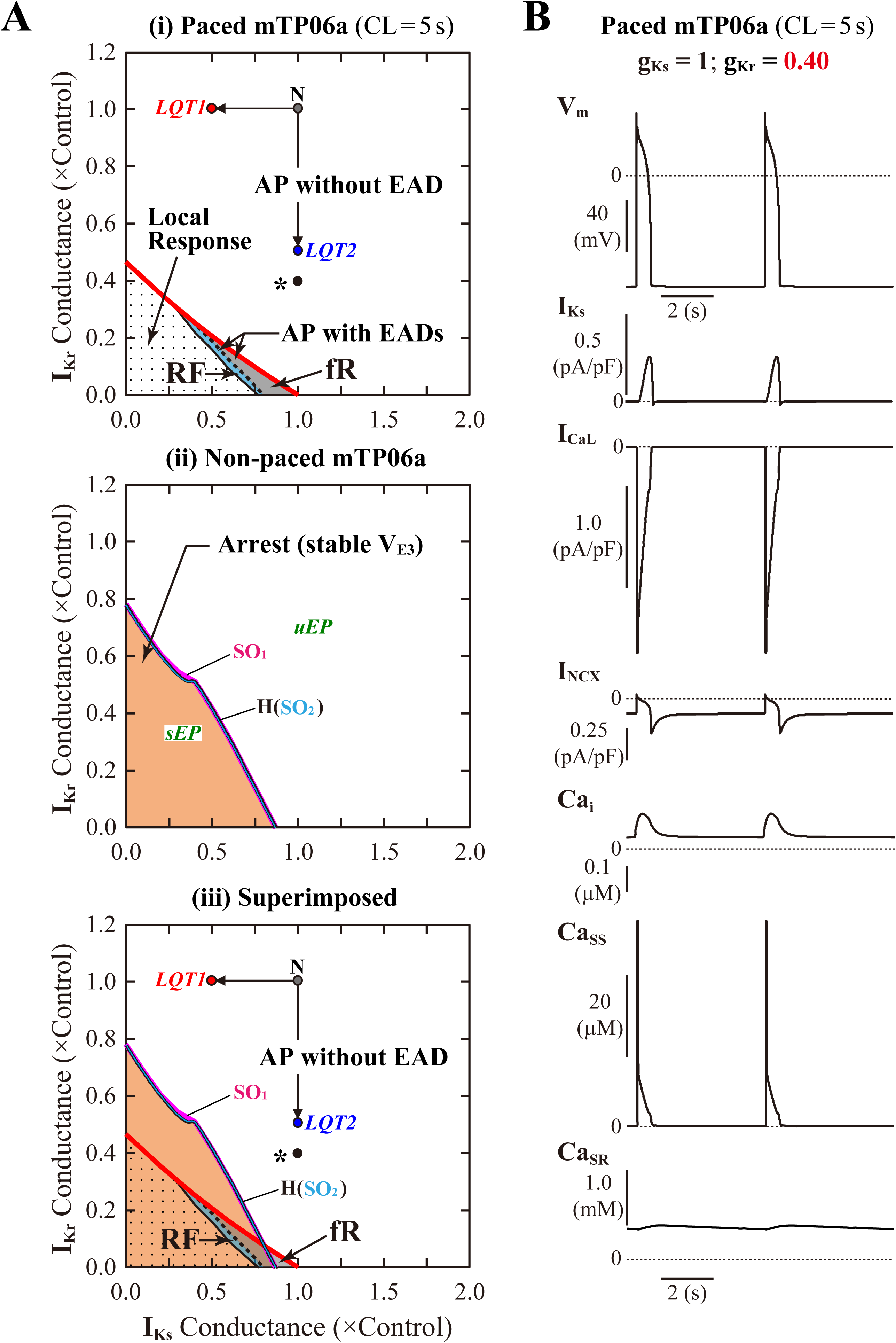
I_Kr_/I_Ks_-dependent EAD generations and bifurcations in the mTP06a model. **(A)** A phase diagram indicating the region of EAD formation (and local responses) in the paced model cell **(i)** and two-parameter bifurcation diagrams for the non-paced model cell **(ii)** on the g_Ks_– g_Kr_ parameter plane. The panel **(iii)** is the diagram for which the phase diagram (i) is superimposed upon the two-parameter bifurcation diagram (ii). In the diagram for the paced model cell (i), the thick red solid, black dashed and thin black solid lines respectively indicate parameter sets of critical points at which short-term EADs (fast repolarization type of APs with EADs), long-term or sustained EADs (repolarization failure type of APs with EADs), and local responses emerged; parameter regions in which short-term EADs, long-term or sustained EADs, and local responses can be observed are shown as the light gray (fR), blue (RF) and dotted regions, respectively. In the diagram for the non-paced model cell (ii), **H**, **SO**_**1**_, and **SO**_**2**_ indicate parameter sets of Hopf bifurcation (HB) points, critical points at which SOs emerged, and critical points at which SOs switched into quiescence, respectively. The parameter region in which convergence to the steady-state (V_E3_) can occur is denoted as the orange region. The labels “*sEP*” and “*uEP*” indicate the areas of stable and unstable equilibrium points (EPs), respectively, divided by the HB curves (H). The points labeled as “N”, “*LQT1*” and “*LQT2*” denote the normal, LQT1, and LQT2 condition, respectively. The asterisks in the panels (i) and (iii) indicate the condition under which the paced model cell behaved as shown in Panel B. **(B)** Simulated behaviors of APs, sarcolemmal ionic currents (I_Ks_, I_CaL_, I_NCX_) and intracellular Ca^2+^ concentrations (Ca_i_, Ca_SS_, Ca_SR_) during 0.2-Hz pacing in the model cell with the parameter set as indicated by the asterisks in Panel A. Temporal behaviors of the variables were computed for 30 min; those for 10 s elicited by additional 2 stimuli are shown.

**Supplementary Figure S3:**
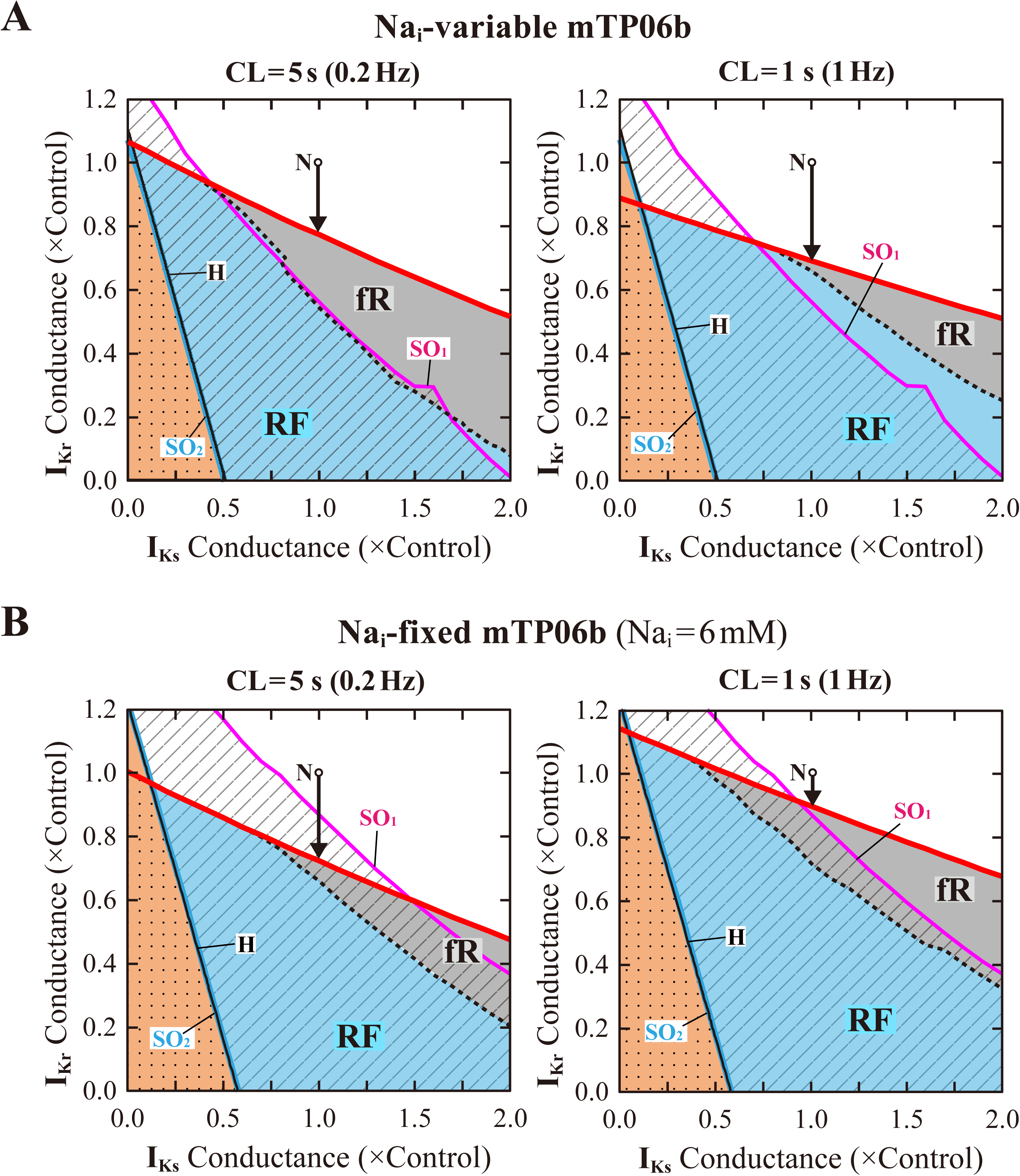
Rate dependence of EAD generation in the mTP06b model. The phase diagrams indicating the region of EAD formation (and local responses) for the paced model cell are superimposed on the two-parameter bifurcation diagrams for the non-paced model cell on the g_Ks_–g_Kr_ parameter plane. The Na_i_-variable **(A)** and Na_i_-fixed **(B)** versions were tested for comparison. The pacing cycle lengths (CLs) were set to 5 s (*left*) and 1 s (*right*). The thick red solid, black dashed and thin black solid lines represent the parameter sets of critical points for occurrences of short-term EADs (APD_90_ < 5 s), long-term or sustained EADs (APD_90_ > 5 s) and local responses, respectively; parameter regions of short-term EADs, long-term or sustained EADs and local responses are shown as the light-gray area labeled as “**fR**” (fast repolarization), blue area labeled as “**RF**” (repolarization failure) and dotted area, respectively. **H**, **SO**_**1**_, and **SO**_**2**_ indicate parameter sets of HB points, critical points at which SOs emerged, and critical points at which SOs switched into quiescence, respectively; parameter regions in which SOs and convergence to the steady state (V_E3_), i.e., arrest, can be observed are shown as the shaded and orange regions, respectively. The points labeled as “N” denote the normal condition.

**Supplementary Figure S4:**
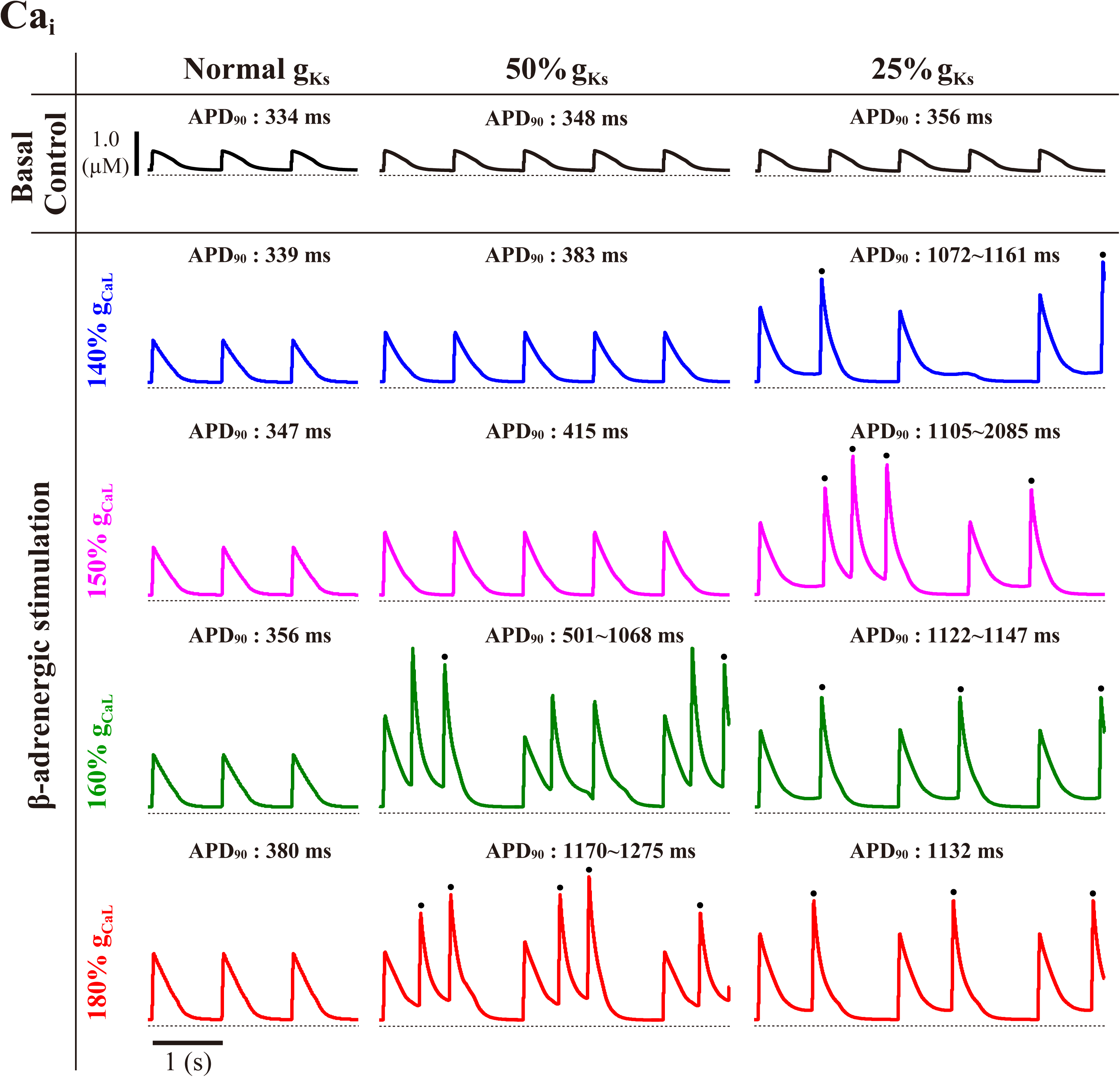
Simulated temporal behaviors of Ca_i_ in the I_Ks_-normal and I_Ks_-reduced (LQT1-type) mTP06b model cells under the basal and β-AS conditions. Model cells were paced at 1 Hz for 30 min under the normal and β-AS conditions as indicated by the points and arrows in **Fig. 5B**. The dots represent Ca_i_ elevations evoked by spontaneous SR Ca^2+^ releases.

**Supplementary Figure S5:**
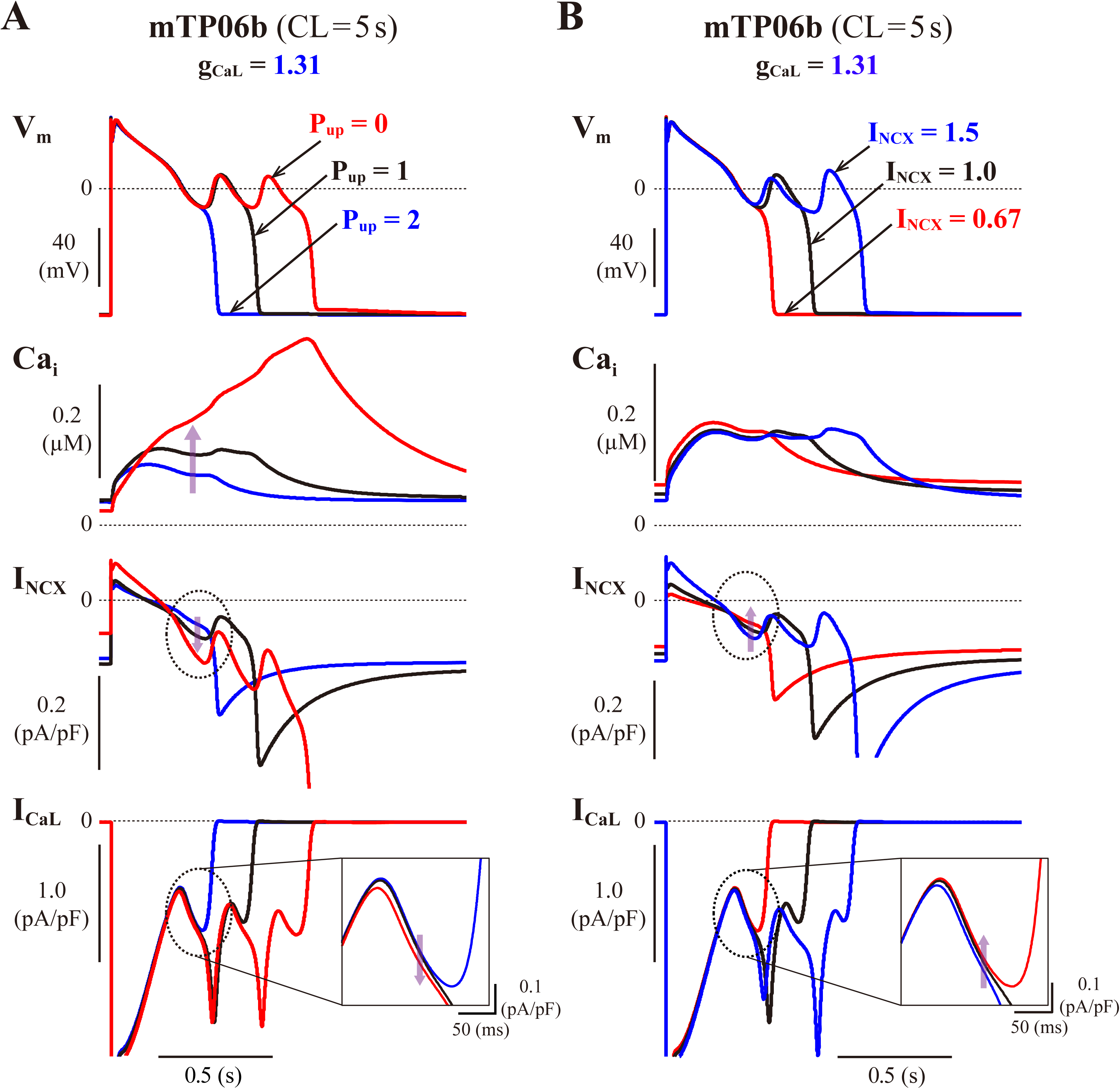
Influences of P_up_ and I_NCX_ on EAD formation in the mTP06b model with an increased g_CaL_. Simulated dynamics of the model cells with various P_up_ **(A)** or I_NCX_ **(B)** values and an increased g_CaL_ (1.31) are shown. P_up_ values were 0 (red), 1.0 (black) and 2.0 (blue) times the control value. I_NCX_ values were 0.67 (red), 1.0 (black) and 1.5 (blue) times the control value. Temporal behaviors of the model cells were computed for 30 min with pacing at 0.2 Hz; the responses to an extra stimulus of V_m_, Ca_i_, I_NCX_, and I_CaL_ for 1.6 s are shown as steady-state dynamics. The dashed ellipses are to focus on the differences in I_NCX_ and I_CaL_ during the preconditioning phase just before initiation of the first EAD. The inset in the I_CaL_ window shows an expanded scale view of I_CaL_ behaviors in the ellipse. The vertical arrows indicate the changes with decreasing P_up_ (A) or I_NCX_ (B).

**Supplementary Figure S6:**
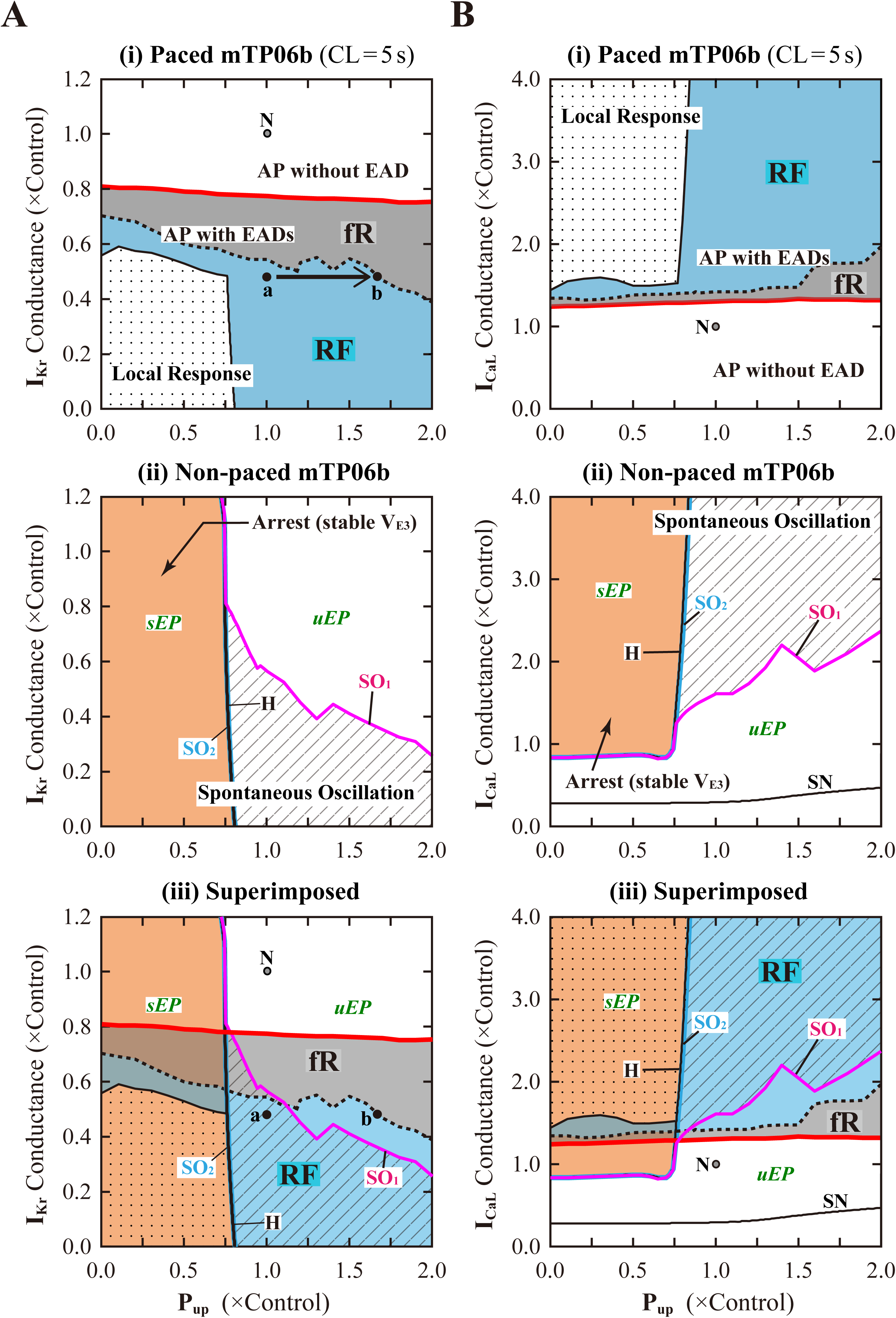
AP behaviors and bifurcations on the P_up_–g_Kr_ **(A)** and P_up_–g_CaL_ **(B)** parameter planes in the mTP06b model. Shown are phase diagrams depicting displacements of critical points for occurrences of short-term EADs (red solid lines), long-term or sustained EADs (black dashed lines) and local responses (black solid lines) in the paced model cell **(i)**, and two-parameter bifurcation diagrams with Hopf (H) and saddle-node (SN) bifurcation curves, and the critical curves for the occurrence (SO_1_) and disappearance (SO_2_) of SOs for the non-paced model cell **(ii)**, as well as the merged diagrams **(iii)** for which the phase diagrams are superimposed on the two-parameter bifurcation diagrams. Representations and symbols are the same as in Fig. 3A and Supplementary Fig. S3.

**Supplementary Figure S7:**
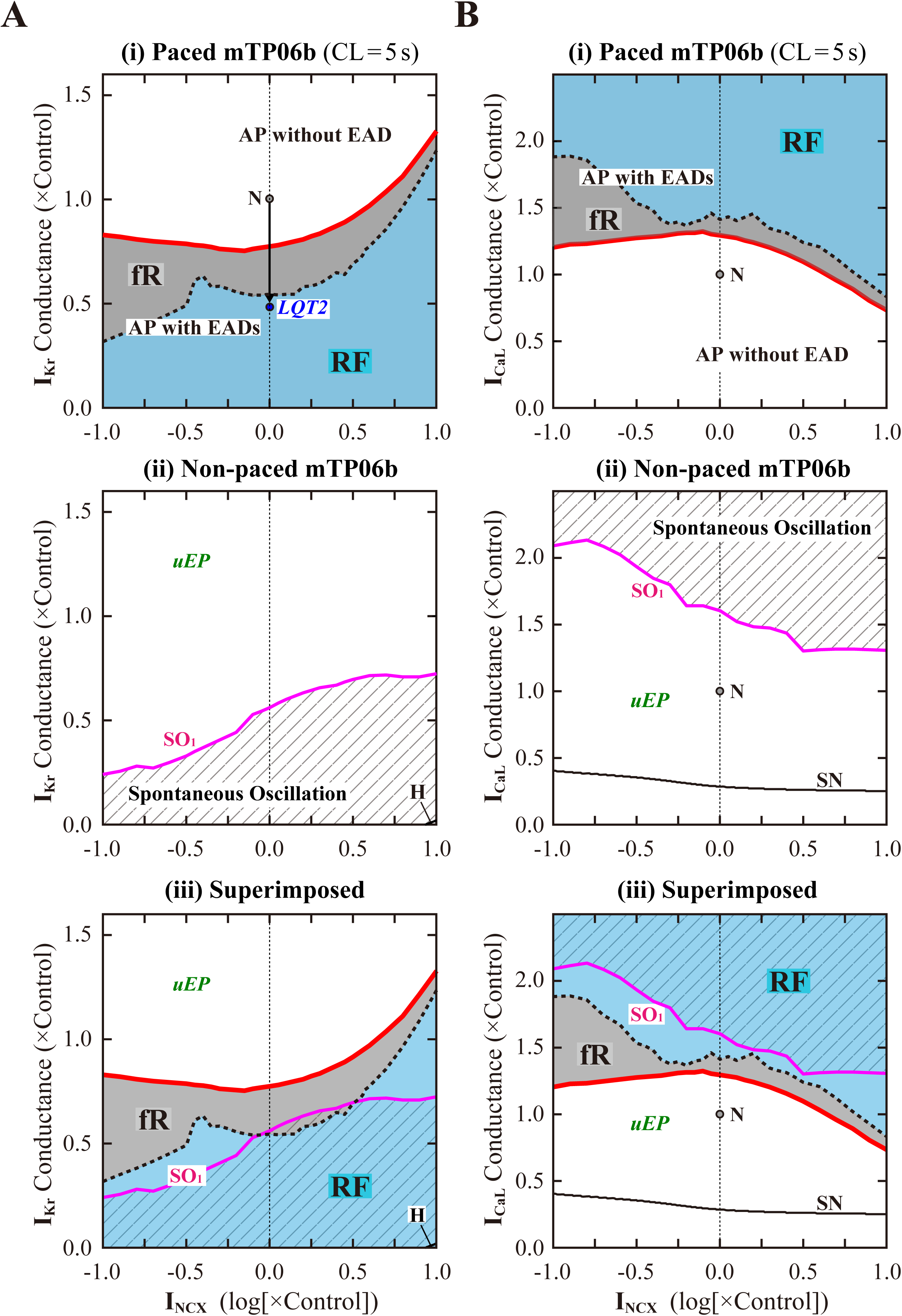
AP behaviors and bifurcations on the I_NCX_–g_Kr_ **(A)** and I_NCX_–g_CaL_ **(B)** parameter planes in the mTP06b model. Shown are phase diagrams depicting displacements of critical points for occurrences of short term EADs (red solid lines) and long-term or sustained EADs (black dashed lines) in the paced model cells **(i)** and two-parameter bifurcation diagrams with the Hopf (H) or saddle-node (SN) bifurcation curve and the critical curve for the occurrence of SOs (SO_1_) for the non-paced model cell **(ii)**, as well as the merged diagrams **(iii)** of the phase diagram (i) superimposed on the two-parameter bifurcation diagram (ii). I_NCX_ density is given as the common logarithm of ratios to the control value. Representations and symbols are the same as in Fig. 3A and Supplementary Fig. S3.

**Supplementary Figure S8:**
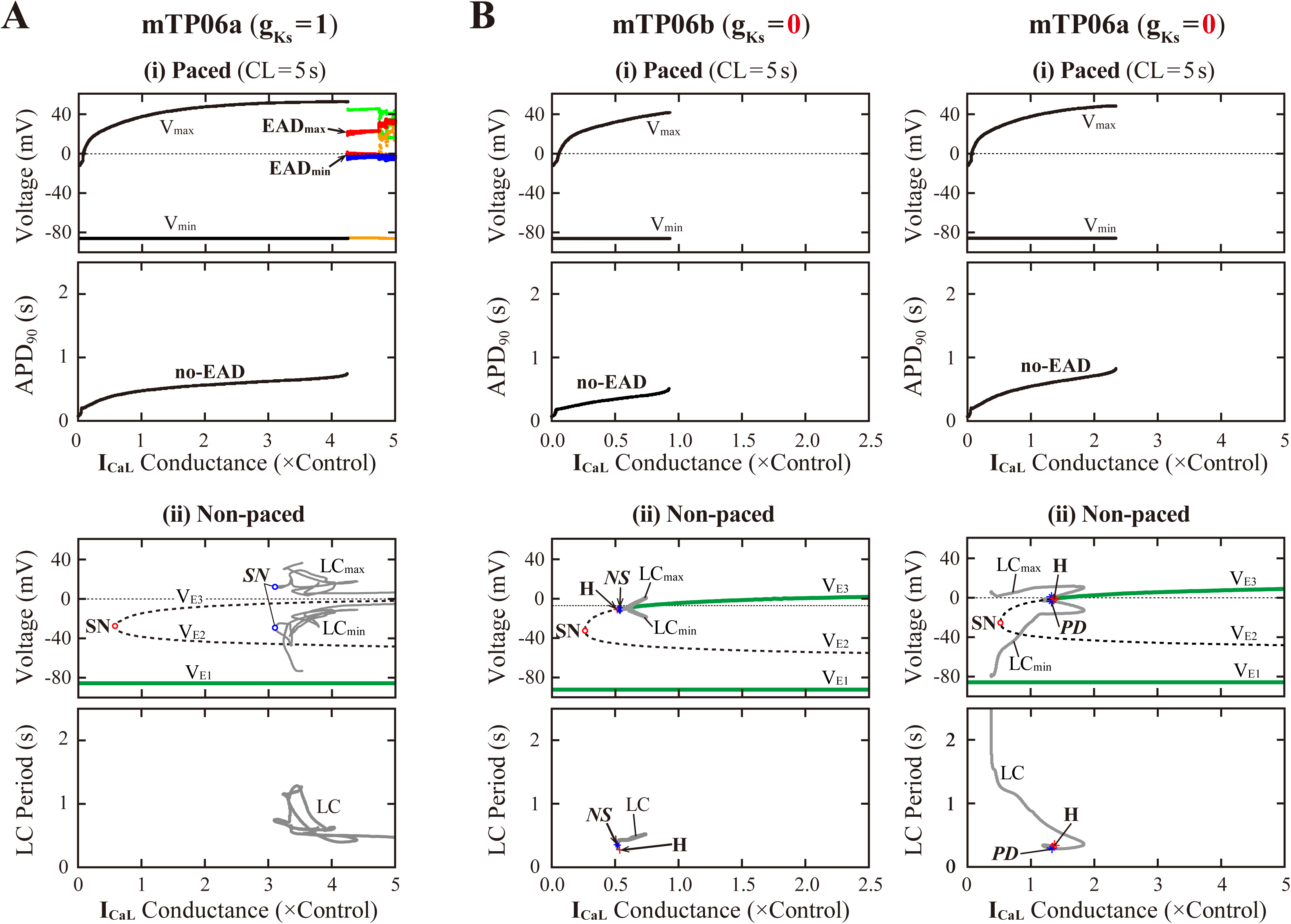
I_CaL_–dependent EAD generations and bifurcations in the I_Ks_-normal and I_Ks_-eliminated mTP06a/b models. Potential extrema of simulated APs and EADs, and APD_90_ are plotted as functions of normalized g_CaL_ for the I_Ks_-normal mTP06a **(A)** and I_Ks_-removed mTP06a/b **(B)** model cells paced at 0.2 Hz **(i)**. AP dynamics during 0.2-Hz pacing were computed for 1 min at each g_CaL_ value, which was increased from 0 to 2.5 or 5.0 at an interval of 0.002. The black dots indicate V_min_ and V_max_ for rhythmic APs, while orange and light green dots in Panel A indicate V_min_ and V_max_, respectively, for arrhythmic APs. EAD_min_ and EAD_max_ were plotted by blue and red dots, respectively. In the diagram for APD_90_, the black dots represent APD_90_ values for regular APs without EAD (no-EAD). One-parameter bifurcation diagrams depicting steady-state V_m_ at EPs (V_E1-3_) and potential extrema of LCs (LC_min/max_), and the period of LCs plotted against g_CaL_ are also shown for the non-paced mTP06a/b model cells **(ii)**. The steady-state branches (V_E1-3_) consist of the stable (green solid lines) and unstable (black dashed lines) segments; the periodic branches are almost always unstable, while stable between the Hopf (H) and Neimark-Sacker (*NS*) or period-doubling (*PD*) bifurcation points.

